# Glutathione metabolism impacts fungal virulence by modulating the redox environment

**DOI:** 10.1101/2024.02.19.581054

**Authors:** Braydon Black, Leandro Buffoni Roque da Silva, Guanggan Hu, Xianya Qu, Daniel F. Q. Smith, Armando Alcázar Magaña, Linda C. Horianopoulos, Mélissa Caza, Rodgoun Attarian, Leonard J. Foster, Arturo Casadevall, James W. Kronstad

**Author notes:** Linda C. Horianopoulos, Wisconsin Energy Institute, University of Wisconsin–Madison, Madison, Wisconsin, 53726, USA. Mélissa Caza, Larissa Yarr Medical Microbiology Laboratory, Kelowna General Hospital, Kelowna, British Columbia, Canada, V1Y 1T2. Rodgoun Attarian, Pfizer Canada, 17300 Trans-Canada Hwy, Kirkland, Québec, Canada, H9J 2M5.

## Abstract

Pathogens must overcome the hostile conditions of their hosts to survive, proliferate and cause disease. The fungal pathogen *Cryptococcus neoformans* is particularly adept at mitigating challenges in the host environment and has developed an arsenal of defense mechanisms to evade oxidative and nitrosative agents released by phagocytic cells during infection. Among these mechanisms, melanin production is crucially linked to both fungal virulence and defense against harmful free radicals that facilitate host innate immunity and clearance of invading pathogens. Here, we employed comparative global metabolomics to demonstrate that metabolism of the antioxidant glutathione (GSH) is inextricably linked to redox-active processes that facilitate melanin production, and that genetic perturbations in GSH biosynthesis affect fungal growth and virulence in a murine model of cryptococcosis. Furthermore, we show that disruption of GSH biosynthesis leads to overaccumulation of reducing and acidic compounds in the extracellular environment of mutant cells. These changes not only impacted melanin formation but also influenced titan cell and urease production as well as survival in macrophages. Overall, these findings highlight the importance of redox homeostasis and metabolic compensation in pathogen adaptation to the host environment and suggest new avenues for antifungal drug development.

---

Evasion of the host immune response is key to the survival and proliferation of microbial pathogens during infection. To facilitate evasion, these pathogens have developed an array of virulence factors that enable them to persist in the host. Such factors include mechanisms for defense against harmful free radicals released by the host during infection as well as the ability to survive at host physiological temperature^1,2^. These traits are particularly relevant in the mammalian host, where innate immunity relies heavily on the generation of reactive oxygen species (ROS) by phagocytic cells to eliminate invading pathogens^1,3^. Thus, processes for maintaining cellular redox homeostasis by acquiring or maintaining intracellular reducing equivalents are crucial for pathogen survival and the spread of disease.

*Cryptococcus neoformans* has emerged as a valuable fungal model for studying such aspects of microbial virulence and host-pathogen interactions^4^. Pulmonary infection by *C. neoformans* begins with rapid proliferation in the lungs followed by dissemination to other organs including the brain^4,5^, which can result in meningoencephalitis that is often fatal^6^. *C. neoformans* evades host immune mechanisms by utilizing several virulence factors which are thought to have evolved in response to environmental predation^5,6^. Notably, melanin is critical for the stress response of *C. neoformans* during infection as it can scavenge harmful free radicals and protect cells from oxidative bursts during phagocytosis^3,7^. Furthermore, cryptococcal meningoencephalitis is associated with fungal-mediated conversion of catecholamines (dopamine, norepinephrine, and epinephrine) from the central nervous system (CNS) into melanin^7^. However, the dependence of melanin on radical polymerization for synthesis has made it difficult to study,^2,8,9^ and the compound requires complex regulatory networks to prevent cell toxicity^10,11^. Thus, mechanisms underlying the redox-mediated processes for melanin production are crucial for understanding virulence and defense strategies of pathogenic fungi, which are becoming more prevalent due to limited diagnostic and treatment capabilities and a continually growing at-risk population^12^. Furthermore, the recent designation of *C. neoformans* as a fungal pathogen of critical importance by the World Health Organization highlights the urgent need to explore the devastating disease caused by this organism – a disease with limited treatment options and no vaccine^12,13^.

Redox-active thiols such as glutathione (GSH) are also key for the oxidative stress response of fungi, and comprise systems that depend on enzymatic and non-enzymatic mechanisms to evade host immunity and cause disease^14–16^. For instance, several GSH-dependent enzymes including glutathione peroxidases and glutathione reductase are important for *C. neoformans* stress response pathways,^17,18^ and others (e.g., glutaredoxins) execute antioxidant functions that require GSH for oxidoreductase activity. For instance, the monothiol glutaredoxin Grx4 regulates iron homeostasis and virulence in *C. neoformans*, and *grx4* null mutants show impaired response to oxidative stress upon iron starvation and/or repletion^19^. GSH is also a signaling molecule for pathogens such as *Listeria monocytogenes*, which requires bacterial and host-derived GSH for activation of the master virulence regulator, PrfA^14,20^. Thus, the importance of GSH is not limited to its redox capabilities. This is clearly demonstrated by the inability of γ-Glu-Cys – a potent antioxidant and direct precursor of GSH – to compensate for non-antioxidant functions in *S. cerevisiae* deletion mutants for Gsh2, the terminal enzyme in the GSH biosynthetic pathway^21^. In fact, GSH is ubiquitous across all kingdoms of life and serves in a vast array of cellular processes; it is also arguably the most important intracellular antioxidant for maintaining redox homeostasis^15,22^. Perhaps unsurprisingly, GSH has also been linked to melanogenesis – a process that is heavily reliant on the cellular redox environment. In human melanoma cells, other mammalian cells, and in birds, depletion of GSH increases melanin deposition^23,24^. High concentrations of GSH also inhibit melanin production, either through direct binding with polyphenol oxidases required for melanin production, or by reduction of radical melanin precursors (e.g., dopaquinone) – with the latter leading to formation of thiol-quinone conjugates used in pheomelanogenesis^25–27^.

GSH is vital for the stress response programs of several microbial pathogens, but its role in the growth, survival, and virulence of *C. neoformans* has not been investigated. In addition to maintaining redox homeostasis during infection, GSH might also support the intracellular conditions needed for melanin production by the fungus – a phenomenon that has not yet been examined in a human fungal pathogen. This study employs genetic and metabolomic approaches to demonstrate that GSH biosynthesis supports the proliferation of *C. neoformans*, and that mutants lacking *GSH2* have attenuated virulence in a murine model of cryptococcosis. We show that loss of Gsh2 perturbs the specific “Goldilocks” redox conditions required for melanogenesis, resulting in an enhanced reducing state that inhibits melanin formation. Furthermore, we demonstrate that an altered redox environment influences titan cell formation, urease production, and interactions with macrophages. Overall, our work demonstrates that *GSH2* and GSH biosynthesis are critical for regulating metabolic and redox-dependent factors that contribute to cryptococcal pathogenesis.

## Results

### Mutants lacking *GSH2* have attenuated virulence

*C. neoformans* requires virulence factors (e.g., capsule, melanin, titan cell formation, and thermotolerance) to evade immune recognition and facilitate disease^5^. We therefore examined the impact of *GSH2* deletion (and thus the lack of GSH) on virulence-related phenotypes using two independent *gsh2Δ* deletion mutants and a *gsh2Δ::GSH2* complemented strain, and found that *gsh2Δ* mutants did not show notable defects in capsule production or heat tolerance (Extended Data Fig. 1a–c). However, the *gsh2Δ* mutants lacked melanin production when grown in medium with L-DOPA (Fig. 1a,b). In fact, *gsh2Δ* mutants were unable to melanize with any of the catecholamine or phenolic substrates tested including L-DOPA, dopamine, epinephrine, or caffeic acid from Niger seed (Extended Data Fig. 1d). Accordingly, we predicted that the virulence of the *gsh2Δ* mutant would be impaired in mice since melanin is linked to virulence and contributes to the neurotropism of *C. neoformans*. Indeed, we found that mice inoculated intranasally with the *gsh2Δ* mutant had reduced disease symptoms and significantly prolonged survival compared to the WT and complemented strains (Fig. 1c,d). In addition, mice infected with the mutant had a lower fungal burden in the lungs, brain, spleen, liver, kidney, and blood at the experimental endpoint (Fig. 1d). Interestingly, fungal burden was particularly diminished in the brain of mutant-infected mice, and it is known that brain tissue is a source of catecholamine precursors for melanin production during infection^7^. Despite multiple attempts using an established protocol, we were unable to isolate melanized cells from tissue homogenates. However, we confirmed that *gsh2Δ* mutants retrieved from lung homogenate could not melanize *ex vivo* (Extended Data Fig. 1e).

**Figure 1.**
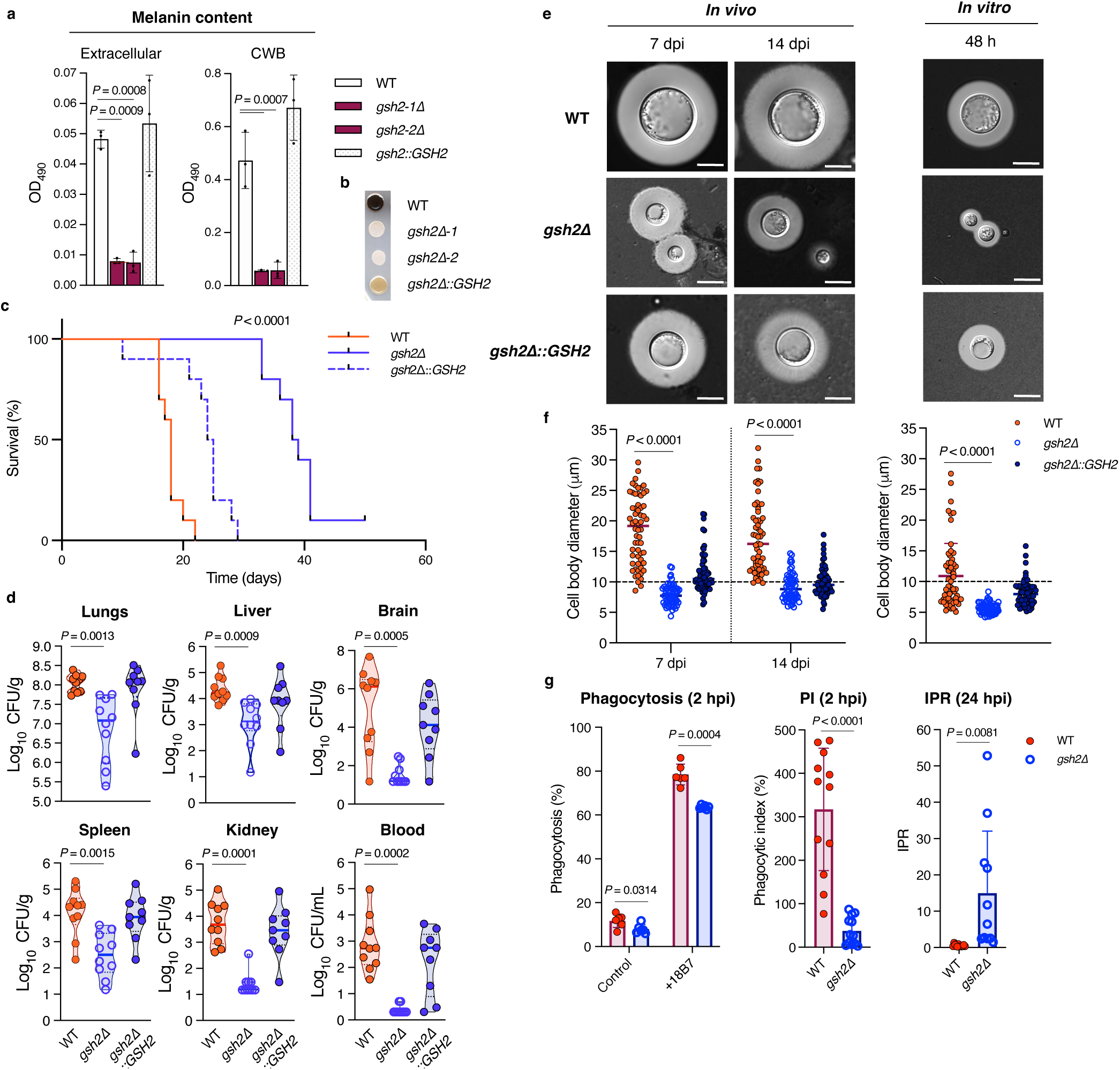
Loss of *GSH2* impairs melanin and titan cell formation and attenuates virulence. **a**, Melanin production of liquid cultures incubated for 48 h in L-DOPA medium. Absorbance of cell supernatant at OD_490_ was measured before (left, extracellular melanin) and after (right, cell wall-bound (CWB) melanin) digestion with 1 M NaOH + 10% DMSO for 1 h at 95^◦^C and normalized by CFU to 10^7^ cells ml^-1^. Bars represent the mean ± s.d. of *n* = 3 biological replicates. Significance relative to WT was calculated using one-way ANOVA with Dunnett’s correction for multiple comparisons. **b,** Melanin production of 10^6^ cells ml^-1^ on solid L-DOPA medium after 48 h incubation at 30**^◦^**C. Images are representative of three biological replicates. **c–d,** Survival curve and measurement of fungal burden in groups of 10 mice after intranasal infection with WT (red), *gsh2Δ* (blue), and *gsh2Δ::GSH2* (blue segmented) strains. Statistical analysis of survival and fungal burden were performed using log-rank (Mantel-Cox) and Kruskal-Wallis (with Dunn’s correction for multiple comparisons) tests, respectively. Fungal burden was measured at the time that the mice were euthanized. **e,** Visualization of fungal cells retrieved from murine lungs (left, *in vivo*) or cell culture (right, *in vitro*) with DIC microscopy. Cells were negatively stained with 0.25 volumes India ink. *In vivo* images are representative of fungal cells retrieved from 8 murine lungs per strain (*n* = 60 cells per sample) at each timepoint (dpi = days post infection), and of three independent experiments per strain (*n* = 50) for *in vitro* images (bars = 10 μm). **f,** Cell body diameters of WT, *gsh2Δ* mutant, and *gsh2Δ::GSH2* strains retrieved from murine lungs (left, *in vivo*) or from cultures grown for 48 h in minimal medium (right, *in vitro*). Measurements represent mean values ± s.d. of 60 cells from each strain per time point with *n* = 8 lungs for *in vivo* measurements and *n* = 3 biological replicates for *in vitro* measurements. **g,** Bone marrow-derived macrophages (BMDM) were isolated from BALB/c mice and infected with WT and *gsh2Δ* mutant strains. Phagocytosis of untreated (control) or opsonized (+18B7) cells was measured after 2 h incubation (left). BMDMs were infected with opsonized *C. neoformans* and lysed after 2 h and 24 h to determine intracellular CFUs, and phagocytic index (PI) was calculated as the CFUs 2 h post-infection (hpi) divided by the total number of macrophages (middle). Intracellular proliferation (IPR) was calculated as the CFUs at 24 hpi divided by CFUs at 2 hpi (right).

The onset of fungal infection and spread to affected organs was also significantly delayed for *gsh2Δ* mutants, which may be partly attributed to an inability of the mutants to form titan cells *in vitro* and *in vivo* – as titan cells are strongly linked to cryptococcal persistence and virulence^4^ (Fig. 1e,f, Extended Data Fig. 1f,g). Additionally, *gsh2Δ* mutants had altered interactions with phagocytic cells including reduced phagocytosis and increased intracellular proliferation compared to the WT (Fig. 1g). Furthermore, mutants had severely diminished proliferation in brain tissue, which was colonized in less than half of mutant-infected mice (Extended Data Fig. 1f). In contrast to the delayed colonization of the brain, fungal burden increased in the lungs, liver, spleen, and kidneys of mice infected with *gsh2Δ* mutants 26 days post-infection, suggesting that *gsh2Δ*-infected mice likely succumbed to cryptococcal pneumonia and/or visceral organ damage rather than brain infection (Extended Data Figs. 1f and 2a). This hypothesis is supported by increases in pro-inflammatory cytokines (IFN-γ, TNF-α, IL-6) observed 26 days post-infection in the lungs of *gsh2Δ*-infected mice, which are absent in these mice 14 days post-infection (Extended Data Fig. 2b). Overall, these findings indicate a role for GSH and the GSH biosynthetic pathway in key aspects of cryptococcal disease including melanogenesis, titan cell formation, interactions with phagocytes, and dissemination to the brain.

### GSH is critical for growth upon nutrient deprivation and is secreted to support the proliferation of GSH-deficient cells

Because GSH is also vital for synthesis of proteins and DNA, we sought to characterize the impact of GSH deficiency in *gsh2Δ* mutants on growth under nutrient-depletion, which mimics the nutrient-sequestered host environment. First, we found that *gsh2Δ* mutants had markedly impaired growth in minimal medium (YNB) over a 72 h period, despite having no growth defect in nutrient-replete media (Fig. 2a,b). Exogenous GSH is also important for combating external stressors (including host stressors during infection^28^). For instance, secretion and extracellular accumulation of GSH in *S. cerevisiae* is crucial for survival and replication under high temperature stress^29^. We therefore investigated whether *C. neoformans* secreted GSH extracellularly – a phenomenon not yet described in this fungus – and if loss of this extracellular GSH pool impacted *gsh2Δ* mutant function. For this experiment, we first quantified extracellular total GSH (reduced plus oxidized GSH) for WT, *gsh2Δ* mutant, and *gsh2Δ::GSH2* complemented strains grown in minimal medium. We found that the WT and *gsh2Δ::GSH2* strains secreted and accumulated GSH extracellularly (Fig. 2c). In contrast, *gsh2Δ* mutants had no detectable extracellular GSH, consistent with the inability of these mutants to synthesize GSH endogenously (Fig. 2c).

**Figure 2.**
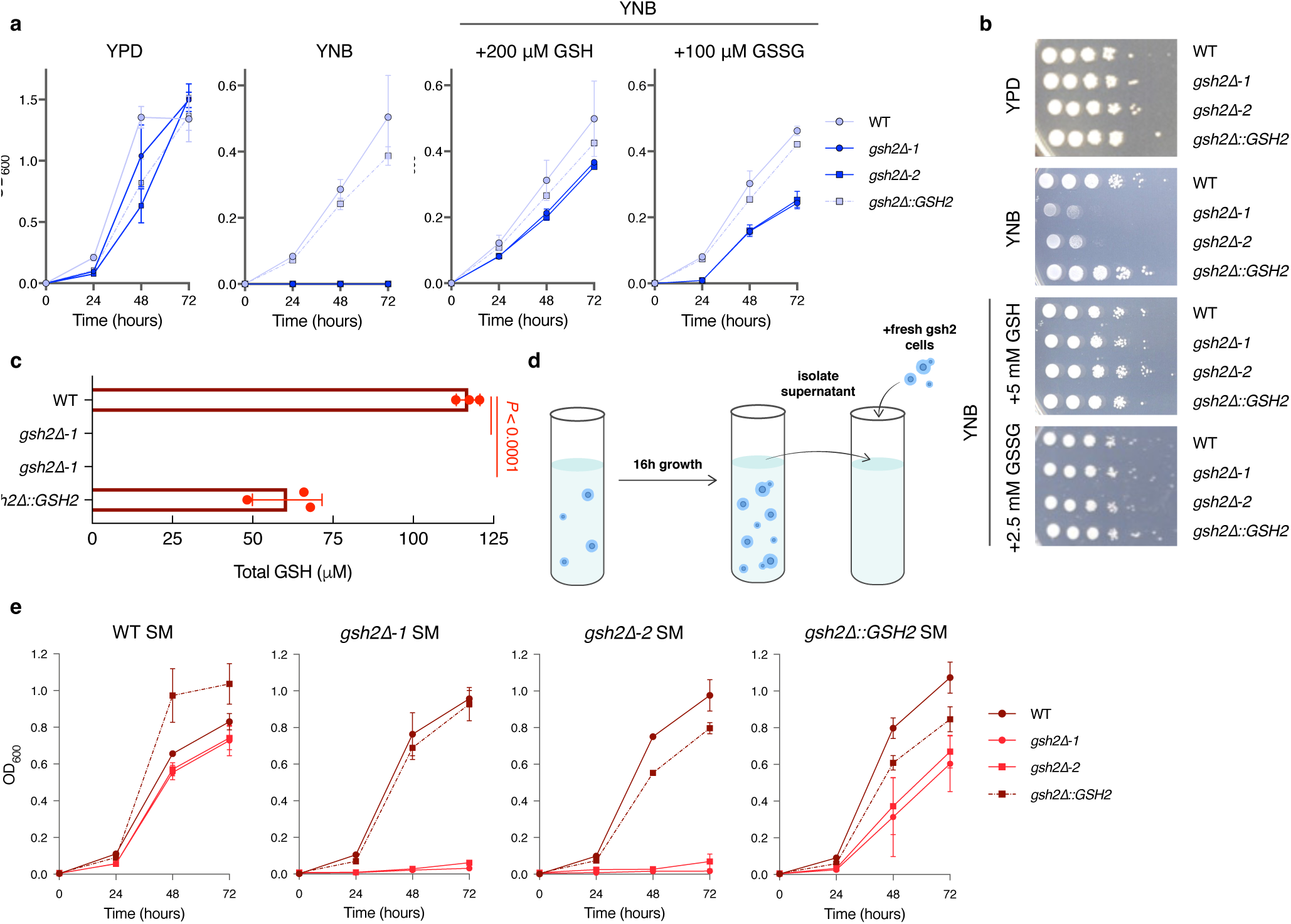
Gsh2 is required for growth upon nutrient depletion and WT cells secrete extracellular GSH. **a**, Growth curve analysis of WT (light blue circles), *gsh2Δ* (blue squares or circles), and *gsh2Δ::GSH2* (light blue squares) strains grown in minimal medium (YNB w/ amino acids plus 2% glucose) with and without GSH supplementation. Data points indicate mean OD_600_ values ± s.d. of *n* = 3 biological replicates at each time point. The initial inoculum for each strain was 2 × 10^4^ cells ml^-1^ in minimal medium and growth was monitored for 72 h, with OD_600_ values measured every 24 h. **b,** Spot assays of growth on solid agar medium starting at 10^6^ cells ml^-1^ with 10-fold serial dilutions with or without GSH or GSSG supplementation at the indicated concentrations. Images are representative of three biological replicates. **c,** Quantification of extracellular total GSH (GSH_t_, reduced and oxidized GSH) for the indicated strains normalized to OD_600_. Bars represent mean values ± s.d. of *n* = 3 independent experiments. Significance was calculated relative to WT using one-way ANOVA with Dunnett’s correction for multiple comparisons. **d,** Schematic for the setup of growth curves in spent minimal medium (SM). **e,** Growth curve analysis of strains growth in SM. SM was isolated from WT (dark red circles), *gsh2Δ* (red squares or circles), and *gsh2Δ::GSH2* (dark red squares) log-phase cells grown in minimal medium for 16 h and was used to monitor growth of fresh cells for each of the strains. Points indicate mean OD_600_ values ± s.d. of *n* = 3 biological replicates at each time point.

Because WT *C. neoformans* secreted and accumulated GSH, we posited that transferring *gsh2Δ* mutants to WT-conditioned medium could elucidate the role of extracellular GSH for GSH-dependent functions in the mutants. Although mutants transferred from an overnight culture in rich medium to minimal medium no longer grew (Fig. 2a), transferring mutants into spent minimal medium (SM) conditioned with WT cells rescued the *gsh2Δ* mutant growth defect to WT levels, despite no additional nutrients being added to the medium (Fig. 2d,e). This outcome suggests that an extracellular factor (likely GSH) supports mutant growth in nutrient limited conditions. Growth of *gsh2Δ* mutants was also partially restored when transferred into SM conditioned with the *gsh2Δ::GSH2* strain (Fig. 2e), although the slower growth relative to cells grown in WT SM may be due to partial complementation and/or lower accumulation of GSH. In contrast, *gsh2Δ* mutants grown in SM conditioned with the *gsh2Δ* mutant did not recover, confirming that mutants are unable to secrete and accumulate extracellular GSH (Fig. 2e).

To eliminate the possibility that other components in the WT SM influenced mutant growth, we tested the impact of exogenous GSH on growth using minimal media (solid and liquid). Indeed, *gsh2Δ* mutant growth in minimal medium was restored to WT levels when supplemented with GSH (Fig. 2a,b). Mutant cultures grown in liquid medium with shaking were particularly sensitive to GSH supplementation and were fully rescued with concentrations as low as 50 μM GSH – a concentration similar to that found in WT supernatant (Fig. 2c, Extended Data Fig. 3a). Intriguingly, addition of the oxidized disulfide GSSG also rescued *gsh2Δ* mutant growth, though to a lesser extent than GSH (Fig. 2a,b). This effect may be attributed to the delay in regenerating GSH via reduction of GSSG by GR^30^, resulting in the initial lag in growth for *gsh2Δ* mutants supplemented with GSSG. Given the overall positive effect of GSH on *gsh2Δ* mutant growth compared to other compounds tested (cysteine, methionine, and ascorbic acid) (Fig. 2 and Extended Data Fig. 3b,c), we conclude that GSH is critical for the growth of *C. neoformans*. This dependence could impact survival and proliferation in vertebrate hosts.

### Loss of *GSH2* does not affect susceptibility to oxidative stress but does impact expression of key antioxidant functions

Because *S. cerevisiae GSH2* deletion mutants accumulate γ-Glu-Cys,^21^ an antioxidant and direct precursor of GSH, we hypothesized that a similar situation would occur in *C. neoformans gsh2Δ* mutants, although this relationship has not yet been demonstrated. Furthermore, we suspected that disruption of GSH biosynthesis could result in accumulation of similar upstream GSH constituents and that loss of extracellular GSH could impact the external redox environment. We therefore tested the sensitivity of *gsh2Δ* mutants to the oxidant H_2_O_2_, and found that mutants treated with H_2_O_2_ had lower ROS accumulation than the WT and complemented strains – a result consistent with the phenotype of *S. cerevisiae gsh2Δ* mutants^21^ (Fig. 3a,b). The observed differences in ROS accumulation between strains led us to investigate potential changes in other antioxidant functions that could potentially compensate for the loss of *GSH2* in the mutants. We found that *gsh2Δ* mutants had enhanced superoxide dismutase (Sod) and catalase (Cat) antioxidant enzyme activities, which are crucial for fungal oxidative stress response and could partly compensate for loss of GSH (Fig. 3c). We further speculated that loss of the extracellular GSH pool in *gsh2Δ* mutants could change the external environment and consequently affect extracellular redox homeostasis. The extracellular composition of the media from mutant cultures was of particular interest, since melanin assembly occurs at the cell wall and is highly dependent on redox-active processes that drive radical polymerization^7,10^. Furthermore, accumulation of L-DOPA in the cell supernatant can cause toxicity and impair growth^31^ – a trend observed in *gsh2Δ* mutants grown in L-DOPA medium, which was rescued with ≥ 50 μM GSH supplementation (Extended Data Fig. 3d,e). We therefore examined the antioxidant/reducing power of the mutant supernatant using an ABTS radical-scavenging assay, which positively correlates reducing capacity with decolorization of a blue/green ABTS radical chromophore. Intriguingly, the supernatant isolated from *gsh2Δ* mutants grown in L-DOPA medium fully quenched the ABTS radical – consistent with the reduced accumulation of ROS in this strain – suggesting a high reducing potential in the extracellular environment (Fig. 3d,e). However, because L-DOPA is a known scavenger of ABTS radicals, we also tested the radical scavenging potential of cells grown in L-asparagine minimal medium (lacking L-DOPA). Both WT and mutant supernatant from cells grown in this medium were unable to quench the ABTS radical, suggesting that accumulated L-DOPA in the *gsh2Δ* mutant supernatant contributed to the ABTS scavenging effect (Extended Data Fig. 3f). However, since *gsh2Δ* mutants had less ROS accumulation in minimal medium than the WT and higher rates of intracellular proliferation in bone marrow derived macrophages (BMDM) (Figs. 1g and 3a,b), we suspected that other mechanisms contributed to the observed changes in redox homeostasis. We therefore postulated that loss of the extracellular GSH pool altered the composition of the mutant supernatant, likely due to accumulation and/or secretion of metabolites upstream of GSH – some of which (e.g., γ-Glu-Cys and cysteine) could compensate for the antioxidant and/or reducing abilities of GSH. In this regard, we did find a modestly elevated thiol content in *gsh2Δ* mutant supernatant relative to the WT or complemented strains (Fig. 3f), and thiols are known to interfere with oxidation of L-DOPA^25–27^. Thus, we surmise that GSH deficiency is sufficient to induce dysregulation of the cellular redox environment and impair formation of virulence-related traits (e.g., melanin and titan cell formation) of *gsh2Δ* mutants.

**Figure 3.**
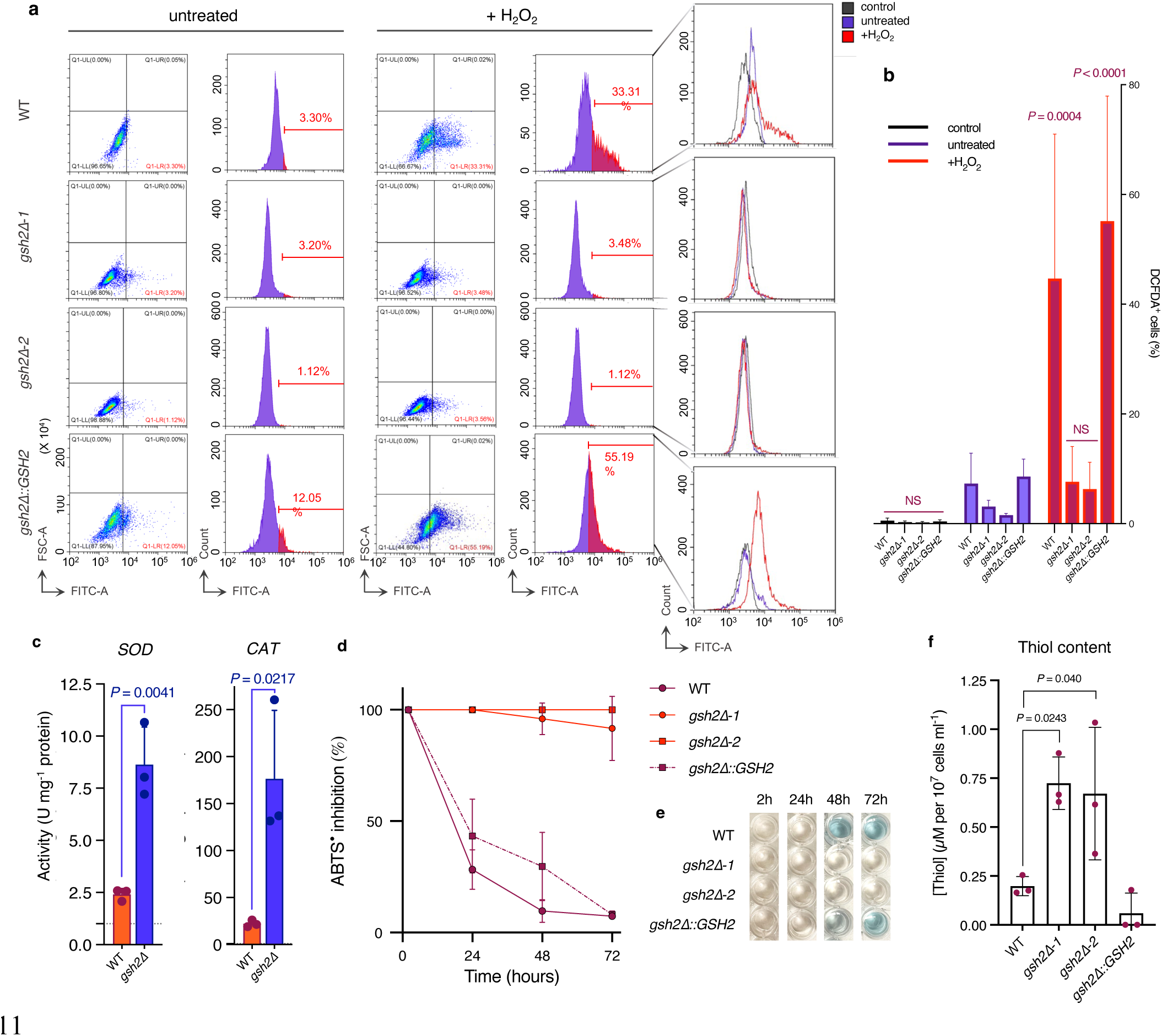
Deletion of *GSH2* reduces susceptibility to H_2_O_2_ stress and alters non-GSH antioxidant functions. **a**, 2D density plots of the indicated strains stained with 16 μM DCFDA (untreated, +H_2_O_2_) or without (control) and analyzed via flow cytometry. Each measurement represents 30,000 gated single cells with or without 1 mM H_2_O_2_ treatment 1 h prior to harvesting from minimal medium. **b,** The percentage of DCFDA-stained (DCFDA+) cells after H_2_O_2_ treatment relative to untreated cells was measured for each strain from **a**. Statistical significance was calculated using a two-way ANOVA with Dunnett’s correction for multiple comparisons. Bars represent mean percentage of DCFDA**^+^**cells ± s.d. **c,** Superoxide dismutase (Sod) and catalase (Cat) enzyme activity (U mg^-1^ protein) of cell lysate normalized by protein concentration. Bars represent the mean ± s.d. and significance was calculated relative to WT using unpaired, two-tailed Student’s *t*-tests. **d–e,** ABTS antioxidant assay for the proportion of ABTS radical (ABTS^•^, blue colouration in **e**) quenched by supernatant isolated from the strains indicated after 72 h incubation in L-DOPA medium. Measurements were taken every 24 h for 72 h and were quantified as a percentage of total quenched ABTS radical (% inhibition). Decreased pigmentation (light blue & clear in **e**) indicates radical scavenging activity. Data points in **c** represent the mean ± s.d. **f,** Fluorescence quantification of supernatant thiol content after 48 h growth in L-DOPA medium for the indicated strains (OD_535_). Bars represent the mean thiol concentration ± s.d. per 10^7^ cells ml^-1^. Data are representative of three biological replicates for each experiment.

### Mutants lacking *GSH2* have dysregulated cellular metabolism

To explain the metabolic changes incited by deletion of *GSH2,* including changes to regulatory machinery for redox control, we characterized and compared the metabolomes of WT and *gsh2Δ* mutant cells grown under melanizing conditions, as melanin synthesis depends on oxidative processes to facilitate polymerization. In particular, we searched for changes in the relative abundance of reducing compounds in culture supernatants that could explain the observed differences in redox potential of WT and *gsh2Δ* mutants. This included quantification of thiol and/or antioxidant compounds directly upstream of Gsh2 (including sulfur-containing compounds and other reductants) with reducing power. Analysis of WT and *gsh2Δ* mutant supernatants and cellular extracts via liquid chromatography-high resolution tandem mass spectrometry (LC-HRMS/MS)-based untargeted metabolomics revealed peak areas of >1500 distinct deconvoluted molecular features (positive ionization mode), of which 439 extracellular features and 229 intracellular features were significantly different (FC > 1.5 or < 0.667, *p* < 0.05) between the WT and *gsh2Δ* mutant strains (Extended Data Fig. 4a). Multivariate analysis using a heat map and principle-component analysis (PCA) revealed a high degree of similarity between WT and mutant cell extracts (Extended Data Fig. 4b,c), but a substantial difference in the relative abundance of metabolites between supernatant fractions of the two strains (Extended Data Fig. 4b,c). Of the 439 differentially abundant features in the supernatant fraction, 294 features had a more-than 1.5-fold increased abundance in the *gsh2Δ* mutant relative to the WT and 80 metabolites had more than a 50-fold increase. However, features with a maximum FC ≥ 50 were interpreted with caution, as excessively high FC values could be attributed to abundances below the limit of quantification in the WT strain.

To evaluate the biological relevance of these metabolic changes, we performed mummichog pathway enrichment analysis (based on KEGG pathway data) and identified significant changes to several key metabolic pathways including amino acid and secondary metabolite metabolism, as well as carbohydrate/energy metabolism (Fig. 4a). We further annotated specific metabolites using a robust scoring technique (see Methods) and confidently identified >100 features in the supernatants and cellular extracts that contributed to the observed pathway changes. We used these annotated metabolites for further analysis (Supplementary Dataset 1). Most strikingly, analysis of the relative abundance of these metabolites showed extracellular accumulation of several aromatic amino acids and weak acids (e.g., dicarboxylic acids, ketones/keto-acids, and hydroxy fatty acids) in the *gsh2Δ* mutant and a significantly depleted intracellular amino acid content (Fig. 4b,c). The mutant also had substantially depleted intracellular levels of key energy pathway intermediates, including adenosine and adenosine monophosphate (AMP), L-carnitine and propionylcarnitine, pantothenate, α-ketoglutaric acid, and ý-nicotinamide adenine dinucleotide (NAD) and nicotinamide (which drive generation of adenosine triphosphate (ATP) in the mitochondria), as well as the urea cycle intermediates ornithine and argininosuccinic acid (Extended Data Fig. 5a). Consistent with the melanin defect and ABTS radical scavenging activity, *gsh2Δ* mutants had a >500-fold increased relative abundance of the melanin precursor L-DOPA (Extended Data Fig. 5b), suggesting that the mutant was unable to utilize L-DOPA for melanin production. Finally, the mutant accumulated significant amounts of extracellular glucosinolates (1,4-dimethoxyglucobrassicin and indolylmethyl-desulfoglucosinolate), sulfur-containing compounds and cysteine derivatives, and other antioxidants that could account for some of the redox changes characterized above (Extended Data Fig. 5). For instance, the antioxidants caffeic acid, salvianolic acid, and pyrogallol-2-O-glucuronide had a more than 50-fold increased abundance in the *gsh2Δ* mutant compared to the WT (Extended Data Fig. 5c,d). Anti-melanogenic compounds were also found, including myo-inositol and hydroxytyrosol, the latter of which was over 150-fold more abundant in the *gsh2Δ* mutant supernatant (Extended Data Fig. 5b). We also detected an approximately 10-fold increase of the cysteine derivative S-(5-histidyl)cysteine sulfoxide, a precursor to the potent antioxidant ovothiol which has Gpx-like activity but has not yet been described in *C. neoformans* (Extended Data Fig. 5c)^32^. Of note, the WT strain had a high intracellular concentration of the antioxidant ergothioneine (EGT), an integral redox buffer in several non-yeast fungi, cyanobacteria, and certain gram-positive bacteria including *Mycobacterium tuberculosis* (Extended Data Fig. 5c)^33^. Though EGT has not been described in *C. neoformans*, our findings further support the proposed interdependency of EGT synthesis on GSH and/or the enzymes involved in GSH biosynthesis^33^.

**Figure 4.**
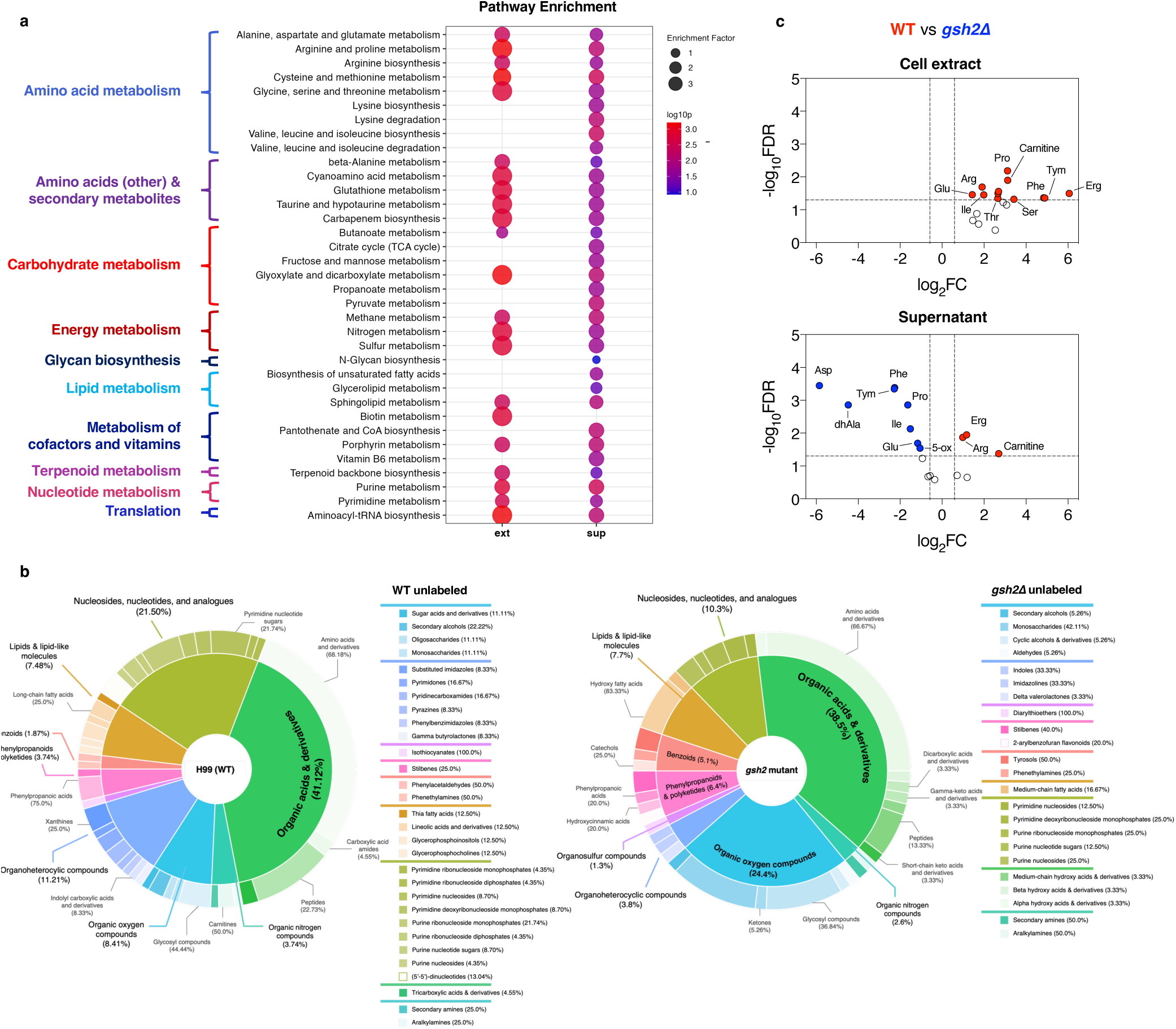
Dysregulation of GSH biosynthesis affects cellular metabolism. **a**, Peak list profile of significantly enriched metabolic pathways using mummichog analysis (v2) from MetaboAnalyst 5.0. Metabolic pathways are grouped by *S. cerevisiae* KEGG pathway module and class datasets. **b,** Classification of differentially abundant metabolites (*p* < 0.05) between *gsh2Δ* mutant (FC < 0.667) and WT (H99) (FC > 1.5) strains using ClassyFire batch compound classification (https://cfb.fiehnlab.ucdavis.edu/). Only metabolites with high confidence annotations (Progenesis QI score ≥ 40) from either positive or negative ionization modes were included in the classification analysis. **c,** Relative abundance of amino acids and derivatives identified via LC-HRMS/MS in both the cell extract (left) and supernatant (right) fractions. The horizontal axis represents directional intensity of the metabolite peak abundance fold change (FC), and the vertical axis represents statistical significance. Thresholds of *p <* 0.05 and FC > 1.5 or < 0.667 (segmented lines) were used to determine significance, which was calculated using an unpaired, two-tailed Student’s *t*-test with FDR correction in MetaboAnalyst (**a, c**). Blue dots = significant metabolites with higher abundance in *gsh2Δ* mutants relative to WT; red dots = significant metabolites with higher abundance in WT relative to *gsh2Δ* mutants.

### Dysregulation of cellular redox homeostasis prevents melanin formation

Our metabolomic analyses predicted several differences between WT and mutant cells, and we examined these alterations using laccase activity and melanin formation as readily assayable phenotypes. Initially, we noted that the abundance of extracellular acids detected by LC-HRMS/MS predicted pH differences between WT and mutant supernatants. We tested this idea and found that supernatant of the *gsh2Δ* mutant culture was more acidic than that of the WT (Fig. 5a). This was true of mutant cells grown in either L-DOPA or L-asparagine minimal media, which demonstrates that accumulated L-DOPA in the mutant supernatant was not responsible for extracellular acidification (Extended Data Fig. 6a). Since *C. neoformans* melanin formation is also influenced by regulation of extracellular pH via the urease-dependent production of ammonia^34^, and mutant growth in L-DOPA medium resulted in low extracellular pH, we investigated urease activity in the mutant. Consistent with an acidic extracellular pH, *gsh2Δ* mutants had significantly impaired urease activity (Fig. 5b). Laccase activity at the cell wall of *gsh2Δ* mutants was also diminished relative to the WT, suggesting that reduced expression and/or improper localization of the enzyme contributed to the melanin defect, potentially due to pH-dependent regulation of laccase activity^35^ (Fig. 5c). Furthermore, buffering the *gsh2Δ* culture medium with 1M MOPS (pH 7.4) restored melanin production, validating an inhibitory role for extracellular acidification in this process (Fig. 5d,e). Metabolomic analyses also predicted changes in antioxidants, which could affect melanin production by inhibiting redox-dependent radical polymerization^7^. Such changes corroborated the potent reducing power of the *gsh2Δ* mutant supernatant (Fig. 3d). We therefore propose that acidification of mutant supernatant (via secretion of acids and impaired urease activity) and changes to extracellular antioxidant composition circumvented the oxidation of L-DOPA, thereby preventing its use in melanin formation by *gsh2Δ* mutants.

**Figure 5.**
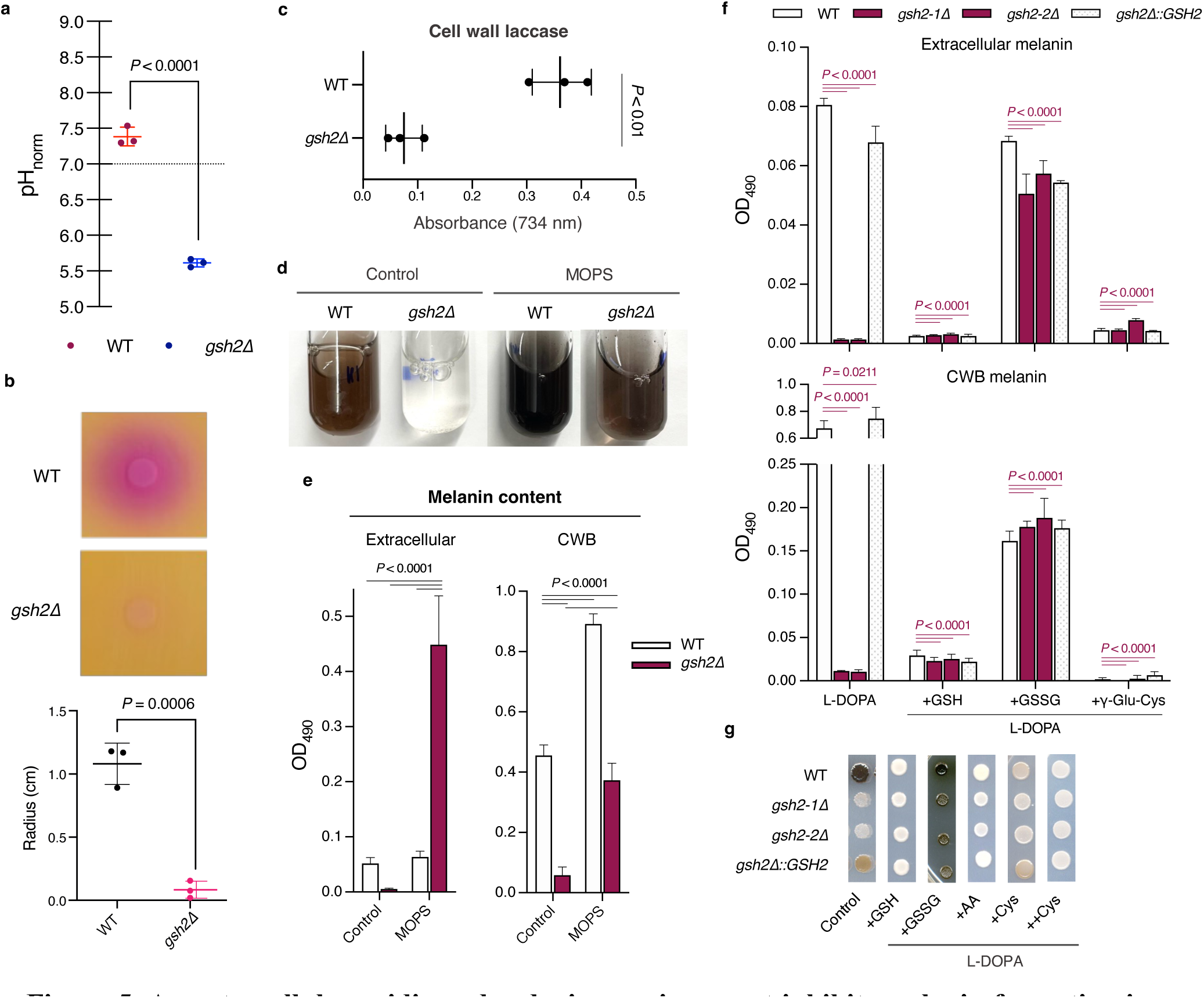
An extracellular acidic and reducing environment inhibits melanin formation in *gsh2Δ* mutants. **a**, pH values of supernatant isolates from WT and *gsh2Δ* mutant cells grown in L-DOPA medium and normalized to 10^7^ cells ml^-1^. **b,** Spot assays of 10^6^ cells ml^-1^ on solid agar urea broth following incubation at 30**^◦^**C for 24 h (left panel). The diffusion radius of ammonia into the medium (pink colouration) indicates urease activity of cells. For **a** and **b** (right panel), statistical significance was calculated using unpaired, two-tailed Student’s *t*-tests, and bars show mean values ± s.d. **c,** Laccase activity of the cell wall fraction of cell lysate measured by conversion of the ABTS substrate, which can be detected at OD_734_. Bars represent mean OD734 values ± s.d. of *n* = 3 biological replicates for each strain. **d–e**, Melanin formation of cells grown in liquid L-DOPA medium with or without 1M MOPS pH-buffering. Extracellular (**d**, left) and CWB (**d**, right) melanin were quantified by measuring absorbance of cell supernatant and digests at OD_490_ (see Methods) and normalizing by CFU to 10^7^ cells ml^-1^. Significance in **d** was calculated relative to WT using one-way ANOVA with Dunnett’s correction for multiple comparisons. Bars represent mean OD_490_ values ± s.d. **f,** Spot assays of overnight cell cultures plated at 10^6^ cells ml^-1^ on agar medium containing L-DOPA to induce melanin production. Mutants lacking *GSH2* showed impaired melanin production on L-DOPA medium and melanin production was not recovered with 5 mM exogenous GSH. The WT and *gsh2Δ::GSH2* complemented strains were also unable to produce melanin in the presence of exogenous GSH (5 mM), ascorbic acid (AA, 10 mM), and 10 mM cysteine (++Cys); melanin production in these strains was substantially inhibited by 5 mM of cysteine (+Cys). Melanin production in *gsh2Δ* mutants was fully recovered upon treatment with 2.5 mM glutathione disulfide (GSSG). Data and images are representative of three biological replicates for each experiment.

To determine whether the impact on melanin formation was specific to loss of GSH or resulted from more general changes to GSH metabolism that impacted cellular redox, we tested other GSH pathway intermediates and ascorbic acid, a well-documented antioxidant. Since GSH, cysteine, and ascorbic acid had comparable radical scavenging activities to the *gsh2Δ* mutant supernatant (Extended Data Fig. 6b,c), we tested the impact of these compounds on melanogenesis of WT and mutant cells. Intriguingly, treatment with GSH, cysteine, or ascorbic acid in high concentrations blocked melanogenesis of all strains including the WT and complement (Fig. 5f,g). We also tested GSH pathway intermediates, as loss of *GSH2* likely impacted the relative abundance of upstream and/or downstream intermediates of GSH metabolism. The thiol-containing GSH precursor γ-Glu-Cys also inhibited melanin production in all strains (Fig. 5f), commensurate with the finding that this compound can compensate for certain antioxidant functions in *gsh2Δ* mutants of *S.* cerevisiae^21^. This compound also fully quenched the ABTS radical chromophore at high concentrations but was less potent than GSH (Extended Data Fig. 6c). We note that γ-Glu-Cys was not detected via LC-HRMS/MS, which may be a result of chemical conjugation and/or utilization by alternate biochemical pathways. For instance, γ-Glu-Cys can directly interact with and neutralize ROS and serves as a cofactor for antioxidant enzymes^36^. In contrast, GSSG fully restored melanin production in *gsh2Δ* mutants and even enhanced melanin production by the WT and complemented strains (Fig. 5f,g). Because *gsh2Δ* mutants retain glutathione reductase (GR) activity, we questioned whether trace amounts of GSH derived from GSSG were responsible for restoring melanin production in GSSG-treated mutants. We therefore performed a GSH titration of mutant cells grown in L-DOPA medium, and found that low levels of GSH (50 – 500 μM) restored extracellular melanin in *gsh2Δ* mutants and further enhanced melanin in the WT (Fig. 6a,b). Notably, the GSH concentrations at which melanin production for *gsh2Δ* mutants was recovered were similar to the amount quantified in the WT supernatant (Fig. 2c). More intriguingly, extracellular melanin production was abruptly blocked in both the *gsh2Δ* mutant and WT when treated with GSH concentrations of 750 μM or higher (Fig. 6a,b), suggesting a ‘tipping point’ for redox-dependent melanogenesis. To our surprise, GSH at all concentrations impaired melanin deposition at the cell wall in the WT and at high concentrations in the *gsh2Δ* mutant (Fig. 6a). However, GSSG (which is oxidizing) and MOPS-buffering restored *gsh2Δ* mutant cell wall-bound melanin – an effect not seen with low-concentration GSH treatment (Figs. 5d–g and 6a). Together, these findings suggest that the synthesis and deposition of melanin depend on the complex regulation of factors influencing both redox and pH.

**Figure 6.**
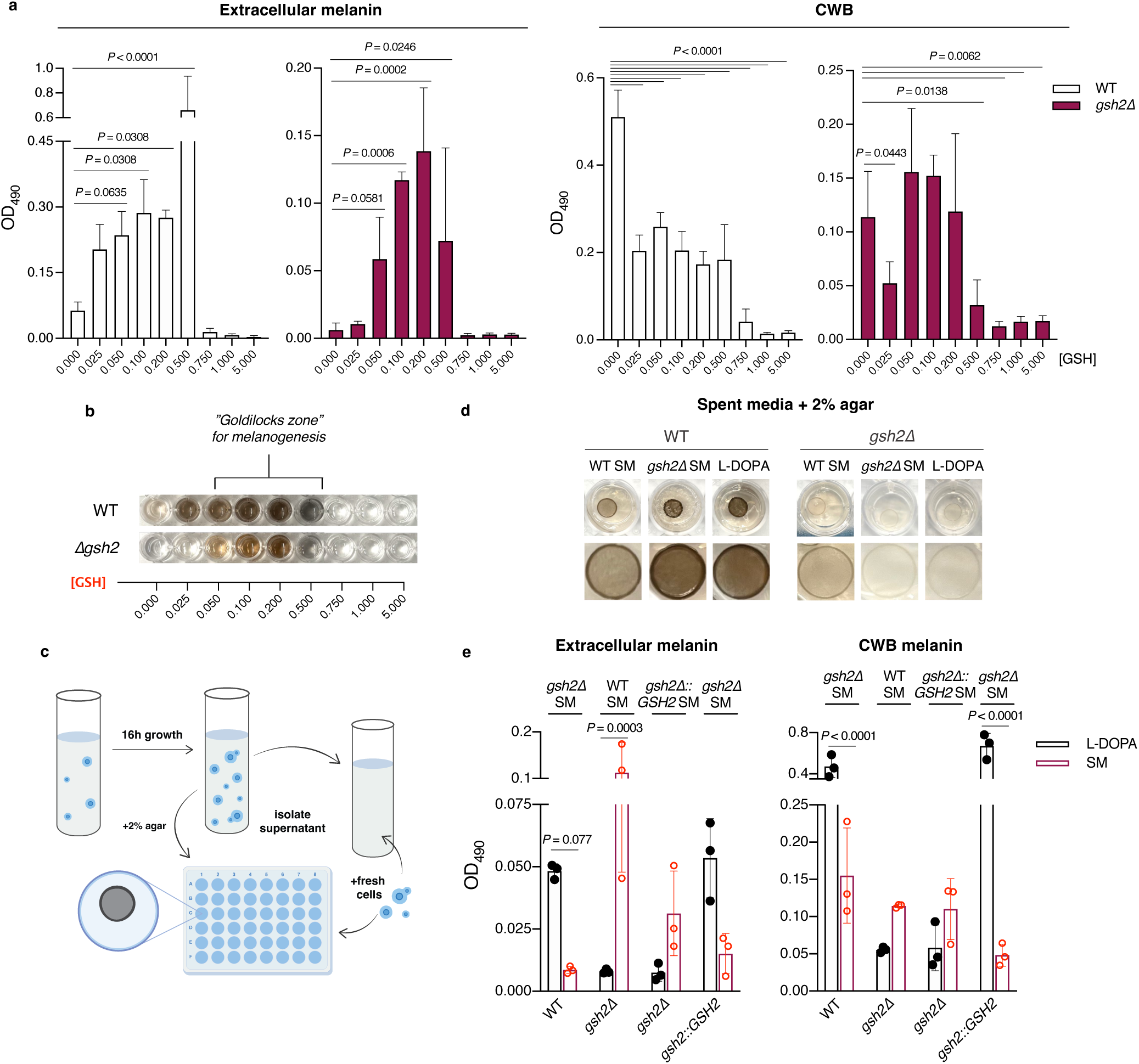
GSH modulates redox homeostasis to influence melanin production. **a**, Melanin formation of cells grown in liquid L-DOPA medium with or without GSH supplementation at the indicated concentrations. Extracellular (**a**, left) and CWB (**a**, right) melanin were quantified by measuring absorbance of cell supernatant and digests at OD_490_ (see Methods) and normalizing by CFU to 10^7^ cells ml^-1^. Significance was calculated relative to cells grown without GSH supplementation using one-way ANOVA with Dunnett’s correction for multiple comparisons. Bars represent mean OD_490_ values ± s.d. **b,** Images of extracellular melanin from WT and *gsh2Δ* mutant cells grown in L-DOPA medium with or without GSH supplementation. **c,** Schematic for the setup of melanization assays in L-DOPA spent medium (SM). **d,** WT and *gsh2Δ* mutant cells melanized for 72 h on agar-supplemented with L-DOPA or L-DOPA SM isolated from log-phase cells grown in L-DOPA medium. Images from the upper panel show individual wells of a 48-well plate and the lower panel shows enhanced-zoom images of spots from the upper panel. **e,** Extracellular and CWB melanin of WT, *gsh2Δ*, and *gsh2Δ::GSH2* cells grown for 48 h in SM isolated from log-phase cells grown in L-DOPA medium. Melanin content was quantified after normalizing by CFU to 10^7^ cells ml^-1^ (see Methods). Bars indicate mean OD_490_ values ± s.d. Data and images are representative of three biological replicates for each experiment.

Since supernatant isolated from the WT had a higher pH, contained low levels of GSH, and restored *gsh2Δ* mutant growth, we reasoned that L-DOPA medium conditioned with WT cells could be used to recover *gsh2Δ* melanin production. We tested this hypothesis and found that mutants transferred to WT SM had significantly increased secreted melanin and modestly elevated cell wall melanin (Fig. 6d,e). In contrast, WT cells transferred to *gsh2Δ* SM had less secreted melanin (*P* = 0.077) and significantly less cell-wall bound melanin (Fig. 6d,e). Taken together, these experiments point to extracellular determinants for melanogenesis and support the conclusion that extracellular changes in *gsh2Δ* mutants lead to a redox environment incompatible with melanin formation.

## Discussion

GSH is a potent antioxidant and key player in the redox defense strategy of many organisms. The compound occurs in naturally high concentrations in most cell types, and is therefore a widespread indicator of the health of the cellular redox environment^37^ – the control of which is critical for normal physiological function and a major determinant of the cellular response to internal and external stressors^1,3^. Here, we demonstrated the regulation of major virulence factors, including melanin and titan cell formation, by manipulation of GSH metabolism and cellular redox homeostasis in *C. neoformans*, a pathogen of global health significance. Our work reveals potential new targets for drug development to combat fungal pathogenesis, and we note that glutathione metabolism has been targeted in human diseases and cancer^38,39^ but not for antifungal drug discovery. Consistent with our study, GSH has been linked to virulence in several fungal and bacterial species and plays a critical role in mitigating the response to host-derived ROS released during the initial stages of infection^15,16,20,28,40^. We discovered a role for GSH metabolism not only in redox control of melanin formation, but also as a systemic regulator of factors that drive growth and virulence. We also demonstrated for the first time that *C. neoformans* secretes and accumulates GSH extracellularly.

The diminished proliferation of *gsh2Δ* mutants – which are non-melanized and have reduced susceptibility to phagocytosis – in the brain of infected mice affirms critical roles for melanin and phagocytic uptake in dissemination (via survival in host macrophages, which can enhance virulence^41^), traversal of the blood-brain barrier, and neurotropism^7^. Perhaps most surprisingly, the *gsh2Δ* mutant failed to produce titan cells *in vitro* and during infection (Fig. 1e,f, Extended Data Fig 1e,f). It is not yet clear whether this defect influences dissemination, though titan cells generate progeny of normal size – some of which are better adapted to host stressors and could more readily disseminate^42^. Furthermore, titan cell morphology can influence immune recognition and skew adaptive immunity to favour cryptococcal dissemination to the CNS^4^. For instance, titan cells appear to induce Th2 immunity which is non-protective and can enhance cryptococcal disease^4,42^. Indeed, we observed an increased abundance of the Th2 cytokine IL-4 in the lungs of WT-infected mice at the time of death, but not in mutant-infected mice (Extended Data Fig. 2b). Conversely, the surge of Th1-associated cytokines (IFN-γ and TNF-α) in *gsh2Δ-*infected mice at the experimental endpoint suggests enhanced inflammation that may damage lungs and exacerbate disease^43^ (Extended Data Fig. 2b). These findings suggest that redox changes due to loss of *GSH2* may generally influence cell morphotypes, impacting both interactions with phagocytic cells and host adaptive immunity. In addition to these defects, we found that loss of GSH in *gsh2Δ* mutants dramatically altered the *C. neoformans* metabolome and resulted in major dysregulation of the cellular redox environment – a key determinant of survival in the host^44^. These findings collectively support GSH metabolism and biosynthesis as potential therapeutic targets for cryptococcosis and provide insights into the mechanisms of cryptococcal disease.

We performed a global metabolomic analysis to identify specific metabolites that might account for altered reducing environment of *gsh2Δ* mutants, and detected multiple potential contributors. For example, the mutant supernatant had substantial quantities of glucosinolates – sulfur- and nitrogen-containing metabolites that contribute to overall antioxidant capacity^45^ (Extended Data Fig. 5c). Mutants also accumulated extracellular phenolic glycosides, including N-acetylserotonin glucuronide and pyrogallol-2-O-glucuronide, which are highly polar compounds that often have strong antioxidant and anticancer activities^46,47^ (Extended Data Fig. 5d). Of note, we found a two-fold increase in pyroglutamic acid, a γ-glutamyl cycle intermediate formed from the GSH precursor γ-Glu-Cys (Extended Data Fig. 5d). This finding is intriguing because human glutathione synthetase deficiency, a heritable amino acid metabolism disorder, also leads to hyperaccumulation of pyroglutamic acid resulting in high anion gap metabolic acidosis – a form of acidosis defined by acid accumulation and low pH of the blood^48^. Consistent with this phenotype, our mutants accumulated and secreted high quantities of amino acids and other weak acids (e.g., lactic acid, dicarboxylic acids, keto acids), which strongly alludes to dysregulated cellular metabolism (Fig. 4c, Extended Data Fig. 5d). The mutants also had significantly lower extracellular pH and urease activity compared to the WT, indicating acidification of cell supernatant (Fig. 5a,b). These finding are corroborated by Mummichog analysis, which showed substantial dysregulation of pathways involved in the metabolism of amino acids/secondary metabolites and glyoxylate/dicarboxylate in both the supernatant and cellular extract fractions (Fig. 4a). Mutant supernatant fractions also contained several amino acid derivatives with known antioxidant and anti-melanogenic properties, including a 150-fold relative increase in hydroxytyrosol^49^. Overall, we propose that accumulation of these compounds (along with excretion of acidic metabolites) leads to redox changes and an enhanced reducing state that is inhibitory for melanin formation.

Despite the oxidative processes involved in melanogenesis, melanin itself is an antioxidant and protects cells against oxidative bursts in the phagosome of phagocytic cells^3,7^. Though melanin has been historically difficult to characterize because of its structural complexity and insolubility, *C. neoformans* is known to secrete urease to modulate extracellular pH and promote melanization^34^ – a process that could influence spontaneous polymerization of radical melanin precursors^2,9^. Thus, the impaired urease and redox activities and low extracellular pH of *gsh2Δ* mutants likely contributed to the lack of melanin in these mutants (Fig. 5a,b). In particular, we suspect that acidification of mutant supernatant precludes chemical oxidation of the catecholamine precursor L-DOPA, as this process is supported by an alkaline pH^50^ and was reversible by MOPS-buffering of *gsh2Δ* mutant media. Indeed, our *gsh2Δ* mutants had a more than 500-fold increase in L-DOPA content relative to the WT, suggesting that spontaneous oxidation of L-DOPA had not occurred (Extended Data Fig. 5b). Low pH and an extracellular reducing environment also support the reduced sensitivity of *gsh2Δ* mutants to H_2_O_2_ stress (Fig. 3). Further, high concentrations of L-DOPA in the mutant supernatant had potent radical scavenging activity and fully quenched the ABTS chromophore monocation – an effect that was comparable to the antioxidants GSH, cysteine, ascorbic acid, and the GSH precursor γ-Glu-Cys (Fig. 3d,e, Extended Data Fig. 6b,c). By comparison, WT cells can utilize L-DOPA for melanin formation and supernatant from the WT was unable to neutralize the ABTS radical (Fig. 3d,e). We therefore hypothesized that the overall perturbations to the *gsh2Δ* extracellular environment were collectively sufficient to block melanogenesis, either by preventing the conversion of L-DOPA into melanin and/or by inhibiting fungal laccase^24,51^. We supported this hypothesis by demonstrating that pH buffering of *gsh2Δ* mutants grown in L-DOPA medium restored melanin formation (Fig. 5d,e). Treatment with exogenous reducing compounds (cysteine, GSH, ascorbic acid, and γ-Glu-Cys) also inhibited melanin formation of the WT and *gsh2Δ::GSH2* strains, and GSSG recovered *gsh2Δ* mutant melanin production (Fig. 5f,g). Since each of the tested reducing agents/thiols blocked melanin production, it is conceivable that accumulation of thiols directly upstream of Gsh2 (γ-Glu-Cys and cysteine) could contribute to the enhanced redox buffering and/or lack of melanin in *gsh2Δ* mutants, likely through diversion to other metabolic pathways. Though γ-Glu-Cys and cysteine were not directly detected by LC-HRMS/MS, several conjugates of these compounds (e.g. pyroglutamic acid, S-(5-histidyl)cysteine sulfoxide) were detected.

Thiols such as GSH are strong antioxidants due to their ability to exist in either thiol or disulfide forms – the former of which can form adducts with or reduce radical melanin intermediates^25,26^. However, these compounds were not identified by LC-HRMS/MS, perhaps due to chemical conjugation (such as thiol-DOPA conjugates), degradation, or association with proteins. We did find an elevated thiol concentration in the *gsh2Δ* mutant supernatant via fluorometric analysis, though the quantity was miniscule and only marginally more abundant than the WT (Fig. 3f). Though specific thiols could not be identified, we observed a substantial abundance of sulfur-containing metabolites in the *gsh2Δ* mutant supernatant which could influence redox homeostasis (Supplementary Dataset 1). Independent of the mechanism, our findings demonstrate an inextricable link between melanogenesis and the status of the cellular redox environment. Specifically, we show that factors affecting extracellular pH can regulate the highly controlled and specific redox conditions required for melanin formation. Such findings support existing evidence that *C. neoformans* secretes extracellular urease to modulate phagosomal pH, which in turn promotes the conditions necessary for melanization, persistence in phagocytic cells, and eventual traversal of the blood-brain barrier^34^. Lack of such regulatory mechanisms, as described with our *gsh2Δ* mutant, resulted in enhanced phagocytosis, lack of melanin, and impaired neurotropism.

Our study sheds light on the complex processes that underlie redox regulation in an epidemiologically important fungal pathogen. Herein, we show that perturbation of the cellular redox environment upsets the specific “Goldilocks” conditions needed for melanin formation, and disrupts the processes required for titan cell formation and dissemination during infection. Specifically, we propose a model wherein deletion of *GSH2* results in systemic dysregulation of cellular metabolism and secretion of metabolites that alter the extracellular reducing environment. These discoveries are consistent with the widespread roles of GSH across fungal and bacterial species^20,52^. However, the link between GSH metabolism and melanin is novel in the context of a human pathogen and warrants further investigation. Thus, our research unveils how GSH metabolism regulates redox control, and provides a strong framework for understanding how microbes mitigate harmful radicals released by host phagocytic cells – a factor strongly correlated with survival and disease dissemination in the host. Finally, our findings can also provide clues to better understand human diseases caused by Gsh2 deficiency and associated disorders of amino acid metabolism that result in anemia and metabolic acidosis^53^.

## Methods

### Strains and growth media

*C. neoformans* var. *grubii* (serotype A) strain H99 was used for generation of *gsh2Δ* deletion mutants and as the wild-type (WT) strain (Table S1). WT and mutant strains were maintained on yeast peptone dextrose medium (YPD, Difco; 1% yeast extract, 2% peptone, 2% dextrose, 2% agar). Cells for growth and phenotypic assays were grown overnight at 30°C (with shaking at 220 rpm) in liquid YPD and were then either harvested or transferred to experiment-specific media for further experimentation. Yeast nitrogen base minimal medium (YNB, Difco; with amino acids, supplemented with 2% glucose, pH 5.6), which lacks GSH, was used as minimal medium. Solid YPD medium containing 200 μg ml^-1^ hygromycin was used for selection of *gsh2Δ* deletion mutants, and medium containing 100 μg ml^-1^ nourseothricin was used to select for the *gsh2Δ::GSH2* reconstituted strain in the *gsh2Δ* mutant background.

### Strain construction

The *gsh2Δ* mutants were constructed with deletion cassette prepared via three-step overlapping PCR using primers listed in Table S2 with WT genomic DNA and the pJAF15 plasmid as templates. The resulting construct was biolistically transformed into the WT strain as described previously^54^, and positive transformants were selected on hygromycin (200 μg ml^-1^) and confirmed via PCR and Southern blot analysis. The *gsh2Δ::GSH2* complemented strain was generated by insertion of a gene encoding a C-terminal Gsh2-HA fusion protein at the native locus. *GSH2*-*HA* was constructed using three-step overlapping PCR with WT and Dnj1-HA genomic DNA as described previously^55^. The resulting construct was biolistically transformed into the *gsh2Δ* mutant background and positive transformants were selected on nourseothricin (100 μg ml^-1^). Primers used for the complement construct are listed in Table S2.

### Murine infection and virulence assays

Virulence of WT, *gsh2Δ* mutant, and *gsh2Δ::GSH2* strains was tested in a murine inhalation model of cryptococcosis using female BALB/c mice (4 – 6 weeks old) from Charles River Laboratories (Ontario, Canada). Briefly, fungal cells were grown in YPD at 30°C overnight, washed twice with PBS, and resuspended in PBS. For survival and endpoint fungal burden, groups of 10 BALB/c mice were intranasally inoculated with a suspension of 2 × 10^5^ cells in 50 μl for each strain and mice reaching the humane endpoint were euthanized by CO_2_ anoxia. For time-course measurements of fungal burden, groups of 8 BALB/c mice were intranasally inoculated with a suspension of 2 × 10^5^ cells in 30 μl for each strain and timepoint. Mice infected for 7, 14, 21, or 26 days were euthanized by CO_2_ anoxia. Mouse organs were harvested by dissection at humane or pre-determined endpoints, weighed, and homogenized in sterile PBS and plated on YPD agar containing chloramphenicol (100 μg ml^-1^) to quantify CFUs for each organ. To enhance experimental rigor and minimize bias for data collection, mice were randomly assigned to experimental groups and chosen at random for euthanasia at pre-determined experimental endpoints. The health status of mice for all experiments was monitored daily post-inoculation. Animal experiments with mice were conducted in accordance with the guidelines of the Canadian Council on Animal Care and approved by the University of British Columbia’s Committee on Animal Care (protocol A21-0105).

### Microscopy of *C. neoformans* and lung tissue sections

For *in vitro* titan cell assays, cells were grown on Sabouraud dextrose agar (SDA; 4% glucose, 1% peptone, 1.5% agar) for 2 – 5 days at RT (∼20–22°C), as described previously^56^. After incubation, 10^7^ cells were resuspended in a T25cm^3^ flask with 10 ml YPD and cultured for 22 h at 30°C with shaking at 150 rpm until stationary phase (approximately 2 × 10^8^ cells ml^-1^). Cells were washed twice with minimal medium, adjusted to 10^6^ cells ml^-1^, and incubated at 30°C for 48 h with shaking at 800 rpm. *In vivo* titan cell formation was evaluated using WT and *gsh2Δ* cells isolated from murine lung homogenate and washed twice with minimal medium. For polysaccharide capsule images, cells were incubated for 48 h in capsule-inducing medium (CIM) prepared as previously described^19,57^. All cells were stained with India ink prior to imaging. Titan cell and polysaccharide capsule formation were evaluated by differential interference contrast (DIC) microscopy using a Zeiss Axiopolan 2 microscope equipped with a Plan-Apochromat 100×/1.46 objective lens and an ORCA-Flash4.0 LT CMOS camera (Hamamatsu Photonics). Zeiss Zen 2 Blue edition (v2.3) and ImageJ^58^ (v2.14.0) were used for collection and analysis of microscopy images, respectively. Cells with a body size >10 μm were considered titan cells, as described previously^56^.

Histological samples were obtained from lung tissue of four infected mice for each strain and timepoint. Lung tissue was stored in 10% formalin and submitted to Wax-It Histology Services Inc. (Vancouver, British Columbia) for processing. Resin-embedded tissues were sectioned, mounted, and stained with hematoxylin and eosin (H&E) prior to imaging.

### *In vitro* macrophage infections with *C. neoformans*

Bone marrow derived macrophages (BMDMs) were isolated from BALB/c mice and differentiated at 37°C with 5% CO_2_ for 7 days in media consisting of DMEM, 10% FBS, 1% nonessential amino acids, 1% penicillin-streptomycin, 1% GlutaMAX (Gibco, 35050061), 1% HEPES buffer, 20% L-929 cell-conditioned supernatant, and 0.1% 2-mercaptoethanol^34^. Differentiated cells were detached from culture dishes using a cell scraper (Falcon, 353085) and washed into DMEM. Cells were seeded at a density of 2 × 10^5^ cells per well in 24-well plates and incubated at 37°C in 5% CO_2_ for 24 h in DMEM supplemented with 100 U ml^-1^ of IFN-γ (Gibco, PMC4031) to activate. Overnight cultures of WT and *gsh2Δ C. neoformans* grown in YPD were washed twice with PBS, counted, and resuspended in DMEM at a density of 2 × 10^7^ cells ml^-1^. Fungal cells were opsonized by incubation with 5 μg ml^-1^ 18B7 monoclonal antibody (mAb) for 1h at RT. BMDMs were then infected with 2 × 10^6^ cells ml^-1^ of opsonized *C. neoformans* and incubated for 2 – 24 h at 37°C. After infection, BMDMs were washed twice with PBS to remove extracellular yeast cells and lysed with 1 ml sterile water. Serial dilutions of BMDM lysate were plated on YPD and grown for 2 – 3 days at 30°C. Colonies were then counted to determine CFUs.

### Lung cytokine isolation and analysis

Lung cytokines were quantified from the supernatant of thawed organ homogenates (stored at - 80°C) from each experimental timepoint using the BD Mouse Th1/Th2/Th17 Cytometric Bead Array (CBA) Kit (BD Biosciences, 560485). Briefly, supernatants were incubated for 2 h at room temperature in equal proportions with anti-cytokine mAb-coated beads and a cytokine phycoerythrin (PE) detection reagent. After incubation, samples were washed once and measured on a Northern Lights full spectrum flow cytometer (Cytek Biosciences). Data were analyzed using FlowJo (BD Biosciences, v10.10) with the CBA Plug-in (v5.2.2).

### Serial dilution spot assays and growth curves

In a typical growth experiment, single colonies of the WT, *gsh2Δ* mutants, or *gsh2Δ::GSH2* strains were selected from an agar plate and incubated at 30°C in YPD medium overnight with shaking at 220 rpm. Cells were harvested and washed twice with sterile water. For spot assays, 10-fold serial dilutions were prepared starting at 10^6^ cells, and 5 μl of cell suspensions were spotted onto solid agar medium. Plates were then incubated at 30°C, 37°C, or 39°C for 2-4 days and imaged. For growth curves, overnight cultures were first counted to determine CFUs. Cells were then diluted in the appropriate medium and grown in either a 96-well plate or 5 ml liquid culture at 30°C for 72 h with shaking at 220 rpm. For YPD and YNB medium growth curves, absorbance at an optical density of 600 nm (OD_600_) was measured every 24 h using a Tecan Infinite M200 PRO microplate reader. For growth in L-DOPA medium, cells were inoculated in chemically-defined L-DOPA medium containing 0.1% L-asparagine, 0.1% dextrose, 3mg ml^-1^ KH_2_PO_4_, 0.25 mg ml^-1^ MgSO_4_·7H_2_O, 1 μg ml^-1^ thiamine, 5 ng ml^-1^ biotin, and 0.2 mg ml^-1^ L-3,4-dihydroxyphenylalanine (L-DOPA; Sigma-Aldrich, D9628) and counted every 24 h to determine CFUs.

### Quantification of melanin production in liquid culture

Cells from overnight cultures were harvested and counted to determine CFUs. Each strain was then diluted and transferred at 1 × 10^6^ cells ml^-1^ to L-DOPA medium with or without supplementation at the concentrations indicated. Cultures were grown for 48 h at 30°C with shaking at 220 rpm, then harvested and counted to determine CFUs. Absorbance of the supernatant was measured at OD_490_ to determine extracellular melanin content. The remaining cell pellet was washed twice with water, diluted to 10^7^ cells ml^-1^, and digested in 100 μl of 1 M NaOH with 10% dimethyl sulfoxide (DMSO) for 1 h at 95°C. Cell digests were centrifuged and the OD_490_ was measured to determine cell wall melanin content. Extracellular melanin content was normalized to growth as determined by CFUs for each strain.

### Media transfer and quantification of extracellular GSH

Following overnight incubation, WT, *gsh2Δ* mutant, and *gsh2Δ::GSH2* cells were washed twice with sterile water, resuspended, and transferred into 25 ml YPD broth at an OD_600_ of 0.3 for an 8 h growth period. Log-phase cells were then harvested, counted to determine CFUs, and diluted in 5 ml of either YNB or L-DOPA medium at concentrations of 3 × 10^4^ or 1 × 10^6^ cells ml^-1^, respectively. After 16 h of growth, cells were harvested via centrifugation and the resulting supernatant was harvested and sterilized using 0.22 μm syringe filter sterilizing units (VWR, 76479-016) to remove residual cells. For YNB medium transfer experiments, fresh overnight cells of each strain were then inoculated into the filtered spent medium (SM) and aliquoted into a 96-well plate at an OD_600_ of 0.0001 and grown for 72 h at 30°C with shaking at 220 rpm and OD_600_ measurements taken every 24 h. For liquid L-DOPA medium transfer experiments, fresh cells were aliquoted into 3 ml spent L-DOPA medium at an initial density of 1 × 10^6^ cells ml^-1^ and grown for 48 h (with shaking at 220 rpm) and then harvested and counted to determine CFUs. Melanin content of liquid cultures was then quantified (as described above). In parallel, fresh cells of each strain were grown in YNB or L-DOPA media to compare cell growth in fresh versus filtered SM. For solid medium experiments, spent L-DOPA medium was supplemented with 2% sterile agar and dispensed into a 48-well plate. After the medium solidified, fresh cells were spotted onto solid spent medium at 1 × 10^6^ cells ml^-1^ and incubated for 48 h at 30°C prior to imaging.

Extracellular GSH was quantified by centrifugation and filter sterilization (VWR, 76479-016) of culture supernatants to remove cells. The concentration of GSH was measured using a glutathione assay kit according to the manufacturer’s instructions (Cayman Chemical, 703002). Absorbance at OD_415_ was measured and GSH concentration was determined by constructing a standard curve using GSH solutions provided by the manufacturer.

### Measurement of ROS content using flow cytometry

Cells were grown for 24 h in liquid YPD medium, washed twice with water and the absorbance at OD_600_ was measured. Resuspended cells were inoculated in fresh YNB (pH 5.6) at an OD_600_ of 0.3 and grown for 16 h with shaking at 30°C. Following a 16 h incubation period, cells were harvested via centrifugation, washed twice using sterile PBS, and resuspended in PBS at a cell density of OD_600_ = 1.0. Cells exposed to oxidative stress were treated with 1 mM H_2_O_2_ for 1 h with shaking at 30°C. After treatment with H_2_O_2_, the treated cells were centrifuged, washed twice, and resuspended in PBS. Cells were then treated with 16 μM of the ROS-sensitive fluorogenic probe 2’,7’-dichlorodihydrofluorescein diacetate (DCFDA, Sigma-Aldrich) for 1 h in the dark at 30°C. Stained cells were then washed once with PBS and ROS levels were analyzed using a CytoFLEX S flow cytometer (Beckman Coulter) with lasers at wavelengths of 405 nm (violet), 488 nm (blue), 561 nm (yellow), and 633 nm (red). Results were gated to single yeast cells and PBS was used as a blank control. Fluorescence of DCFDA was measured using the FITC-GFP channel, and data were acquired and analyzed using the CytExpert cytometry analysis software (Beckman Coulter).

### ABTS free-radical scavenging, thiol quantification, and antioxidant enzyme activity assays

Preparation of the radical ABTS solution was performed as previously described^59^. To examine the radical scavenging activity in WT and *gsh2Δ* mutant culture supernatants, cells were first grown at 1 × 10^6^ cells ml^-1^ in L-DOPA medium, centrifuged and 20,627 × *g*, and 1 ml of the supernatant was collected and dried overnight at room temperature via rotary evaporation. Samples were then resuspended in 200 μl of nanopure water, and 10 μl of each sample was transferred to a 96-well plate. For testing specific chemicals, 10 μl of each compound was serially diluted to the indicated concentrations. Then 190 μl of radical ABTS solution was added to each sample, as described previously^59^. Following a 5 min incubation in the dark, absorbance at OD_734_ was measured to quantify decolourization.

Thiol concentration of culture supernatants was measured using a fluorometric thiol assay kit according to the manufacturer’s instructions (Sigma-Aldrich, MAK151). Fluorescence was measured with a BioTek Synergy H1 microplate reader (Agilent) at an excitation wavelength of 490 nm and an emission wavelength of 525 nm. Thiol concentration was determined by constructing a standard curve using serial dilutions of a GSH standard provided by the manufacturer and normalized by CFU.

The activities of superoxide dismutase and catalase were measured using enzyme-specific colorimetric assay kits according to the manufacturer’s instructions (Cayman Chemical, 706002 and 707002, respectively). Absorbance was measured at 440 nm (Sod) or 540 nm (Cat) using a Tecan Infinite M200 PRO microplate reader, and enzyme activity was normalized to protein concentration as determined by Pierce BCA protein quantification (Thermo Fisher Scientific, 23225) relative to a BSA standard.

### Measurement of urease and cell wall laccase activity

Overnight cultures were harvested and washed twice with PBS at 4,000 × *g* for 4 min. For urease assays, cells were diluted 1:40 in PBS and 25 µl of diluted culture was spotted onto urease detection agar medium (2% urea, 1.5% agar, 0.2% monopotassium phosphate, 0.1% peptone, 0.1% dextrose, 0.5% sodium chloride, 0.0012% pH 6.8 phenol red). Plates were incubated for 24 h at 30°C and scanned using a CanoScan 9000F at 600 DPI. Pink halos indicative of urease activity were measured using ImageJ software^58^. For cell wall laccase activity, washed cultures were added 1:100 in 100 ml of L-asparagine medium (0.1% L-asparagine, 0.1% dextrose, 3mg ml^-1^ KH_2_PO_4_, 0.25 mg ml^-1^ MgSO_4_·7H_2_O, 1 mg ml^-1^ thiamine, 5 ng ml^-1^ biotin) and grown for 48 h, shaking at 120 rpm. Following incubation, cultures were harvested and washed twice in PBS at 14,000 × *g* for 15 min. Cells were then resuspended in 10 ml of cold PBS and incubated with cOmplete mini EDTA-free protease inhibitor (Roche, 11836170001) for 30 min at 4°C. Cells were kept on ice and lysed using a French pressure cell press (Glen Mills) with 5 passes at 25,000 psi. The resulting lysate was then centrifuged at 3,000 × *g* for 10 min and the supernatant was removed. The resulting cell wall pellet was washed twice in PBS at 3,000 × *g* for 10 min and resuspended in 10 ml PBS. Then, 180 µl aliquots of resuspended cell pellet were added to a 96-well plate in triplicate and 20 µl of 10 mM ABTS was added to each sample. Plates were incubated for 2 h at 30°C with constant shaking. Absorbance at 734 nm was measured after 2 h using a SpectraMax iD5 spectrophotometer.

### Cell preparation for metabolome profiling

Samples for metabolite analysis were incubated for 48 h in L-DOPA medium starting at 1 × 10^6^ cells ml^-1^. WT and *gsh2Δ* mutant cells were then harvested via centrifugation at 15,493 × *g* for 10 minutes at 4°C and separated into supernatant and cell pellet fractions. Both fractions were kept on ice. Next, 2.5 volumes of 100% MeOH + 0.05% butylated hydroxytoluene (BHT; Cayman Chemical) was added to the supernatant fraction to limit oxidation and/or isomerization of reactive compounds^60^. The resulting mixture was incubated for 1 h at -20°C to precipitate residual salts from the chemically defined medium. Supernatant fractions were then centrifuged (20,627 × *g* for 10 minutes, 4°C) and a 1 ml aliquot was removed and stored on ice. Two-phase extraction was then performed as described previously, with slight modifications, to isolate intracellular contents of fungal cells^61^. First, the remaining cell pellets were washed twice with pre-cooled nanopure water and 150 μl 100% MeOH + 0.05% BHT was added. The resulting solution was mixed by vortexing and then shaken gently for 2 min at 4°C. Pre-cooled nanopure water (200 μl) was added with ∼150 μl glass beads. Samples were homogenized using a Mini-BeadBeater-8 (BioSpec) for three rounds of 20 s each, with 30 s resting on ice between each round. A 500 μl aliquot of methyl-tert-butyl-ether (MTBE) was then added to each sample followed by vortexing to mix and gentle shaking for 10 min at 4°C. After centrifugation (20,627 × *g* for 10 minutes, 4°C) to induce phase separation, the aqueous layer was extracted and placed in a separate microfuge tube. Both the aqueous cell extract and cell supernatant were then dried using a rotary evaporator overnight at room temperature. Fractions were stored at -80°C until LC-HRMS/MS analysis.

### Liquid chromatography-mass spectrometry (LC-HRMS/MS)

For untargeted metabolomics analysis, samples were analyzed using an Impact^TM^ II high-resolution mass spectrometer from Bruker Daltonics (Bruker Daltonics, Bremen, Germany) coupled with an Elute UHPLC system (Bruker). Separation of compounds was achieved using a multigradient method on an Inertsil Ph-3 UHPLC column (2 µm, 150 x 2.1 mm) (GL Sciences) equipped with a Ph-3 guard column (2 µm, 2.1 x 10 mm). The mobile phase consisted of water (A) with 0.1% v/v formic acid and methanol (B) with 0.1% v/v formic acid. The separation was conducted using a multi-step gradient ranging from 5% to 99% mobile phase B over 18 minutes as follows: 0 min 5% B; 0–1 min 5% B; 1–8 min 35% B; 8–10.5 min 99% B; 10.5–14 min 99% B; 14–14.5 min 5% B; 14.5–18 min 5% B. The column temperature was set to 55°C, while the autosampler was maintained at 4°C, and the flow rate was 0.3 ml min^-1^.

Aliquots of 100 µL from each sample were pooled to generate a quality control sample (QC) used for evaluating instrument performance. Quality control sample was injected every six samples. Data-dependent acquisitions were conducted in positive (ESI+) and negative (ESI-) ionization modes to obtain precursor and fragment ion information for annotating compounds. For ESI+, the mass spectrometer settings were as follows: capillary voltage of 4500 V, nebulizer gas pressure of 2.0 bar, dry gas flow rate of 10 L min^-1^, dry gas temperature of 220°C, mass scan range of 50-1300 m/z, spectra acquisition rate of 10 Hz, and cycle time of 0.4 s. Collision energy of 20 V was ramped through each MS/MS scan from 100 to 250%. For ESI-, the capillary voltage was set at -3500 V. The mass spectrometer was calibrated with sodium formate at the beginning of each run to ensure accuracy. Average mass error in annotation was below 2.0 ppm (Supplementary Dataset 1).

### Data processing and annotation of metabolites

Raw data processing was performed using Progenesis QI™ software (V3.0.7600.27622) with METLIN^TM^ plugin V1.0.7642.33805 (NonLinear Dynamics) and entailed peak picking, alignment, normalization, and database searching^62^. Annotations were performed as previously described^62,63^. However, to increase confidence in annotations, only experimental MS/MS data was used for querying and matching against the *in-house* library (Mass Spectrometry Metabolite Library of Standards, MSMLS, supplied by IROA Technologies), METLIN^TM^ and MassBank of North America^64,65^. A Progenesis QI score ≥ 40 was considered to select a candidate for annotation in accordance with reporting criteria for chemical analysis suggested by the Metabolomics Standards Initiative (MSI)^66,67^. Ions generated from QC samples were retained for annotation and included in the dataset if the coefficient of variation (CV) for abundance did not exceed 25%. When compounds were detected in both ion modes, the one with the highest signal-to-noise ratio was retained. Relative quantities of metabolites were determined by calculating their corresponding peak areas. Finally, putative metabolites with high confidence annotations were categorized using the ontology-based ClassyFire Batch Compound Classification web platform (https://cfb.fiehnlab.ucdavis.edu/).

Data was normalized using a robust Progenesis QI^TM^ built-in approach designed explicitly for untargeted metabolomics^62,63,68^. Progenesis QI^TM^ employed median and means absolute deviation analysis based on all detected abundances to reduce bias and noise in the data^69^. A unique gain factor is then calculated for each sample (represented by a scalar multiplier αk) and compound ion abundances were adjusted to a normalization reference run^68^. In brief, all compounds were included to normalize the data, resulting in an aggregate matrix with measurements for every compound ion in each run. This aggregate matrix was used to generate a ratio for comparison of compound ion abundance in a particular run with the corresponding value in the normalization reference (typically a QC). Ratios were log transformed to create a normal distribution for all ratio data within each run for all samples. Finally, scalar estimations were employed to align log distributions with that of the normalization reference. The tight clustering exhibited by quality control (QC) samples at the center of the PCA analysis indicated excellent system performance and stability (Fig 4C).

### Quantification and statistical analysis

Statistical analyses were performed using Prism 8 (GraphPad Software), unless otherwise specified. Statistical significance between two groups was calculated using unpaired, two-tailed Student’s *t*-tests; significance of three or more groups was calculated using one- or two-way analysis of variance (ANOVA) and corrected for multiple comparisons using the methods indicated.

Metabolomics data was analyzed using the web-based platform Metaboanalyst 5.0 (https://www.metaboanalyst.ca/faces/home.xhtml). Univariate analysis of peak intensity values was performed by volcano plot analysis wherein *p-*values were determined with a two-sided Wilcoxon rank-sum test and then adjusted for multiple testing by FDR correction using an adjusted *p*-value (*q*) threshold of *q* = 0.05. Adjusted *p*-values were plotted on a negative log_10_ scale. Volcano plot fold-change (FC) values were calculated as the ratio of peak intensity means between samples for each feature and plotted on a log_2_ scale with cut-off values of FC > 1.5 or < 0.667.

Multivariate analysis was performed to identify interactions and changes in the overall metabolome profile of WT and *gsh2Δ* mutant strains, as well as supernatant and cellular extract fractions. MetaboAnalyst was used to conduct a Principal Component Analysis (PCA) and a heat-map analysis to visualize distribution of samples and relative intensity of LC-HRMS/MS features, respectively. Metabolic pathway enrichment analysis was performed using MetaboAnalyst peak list profiling which utilizes the mummichog v2 algorithm, based on KEGG pathway data, to quantify enrichment of putatively annotated peaks at the network level, as described by Li et al.^70^. The *S. cerevisiae* KEGG pathway data was used for enrichment analysis as this dataset most closely resembles *C. neoformans* pathway information of datasets available on the MetaboAnalyst platform.

## Data availability

The authors declare that all data supporting the findings of this study are available within the main text and Supplementary Information. Source data are provided with the paper.

## Acknowledgements

The authors thank Dr. Tao Huan for useful discussions. Research reported in this publication was supported by the National Institute of Allergy and Infectious Diseases of the National Institutes of Health under award number R01AI053721 (to J.W.K.). A.C. was supported in part by the National Institutes of Health (NIH) grants AI052733, AI15207, AI171093-01 and HL059842. The content is solely the responsibility of the authors and does not necessarily represent the official views of the National Institutes of Health. Additional support came from a UBC Four Year Doctoral Fellowship (to B.B.), and a Natural Sciences and Engineering Research Council of Canada (NSERC) Postgraduate Scholarship – Doctoral (to B.B.). L.C.H. is an NSERC postdoctoral fellow. J.W.K. is a Burroughs Wellcome Fund Scholar in Molecular Pathogenic Mycology, and the Power Corporation Fellow in the Canadian Institute for Advanced Research (CIFAR) Program: Fungal Kingdom, Threats & Opportunities. Untargeted metabolomic analysis was performed in the Life Sciences Institute (LSI) Metabolomics Core Facility of the LSI at UBC, supported by the Canada Foundation for Innovation, the BC Knowledge Development Fund, the LSI, and the UBC GREx Biological Resilience Initiative.

## Author contributions

B.B. and J.W.K. conceptualized this project. B.B. developed methodology with guidance from J.W.K., A.C., and D.F.Q.S. and performed all growth experiments, flow cytometry, melanin/ABTS and colorimetric/fluorometric kit assays, and data analysis. B.B. wrote the manuscript and J.W.K. provided edits. All authors participated in the review and editing of the manuscript. G.H., L.B.R.S., X.Q., B.B., L. C. H. and M.C. performed animal work and X.Q. imaged cells. Macrophage experiments were conducted by L.B.R.S., X.Q., and B.B., and L.B.R.S. performed all cytokine profiling experiments. G.H., L.C.H. and R.A. designed and constructed the mutant and complemented strains used in this study. A.A.M. conducted all LC-HRMS/MS experiments with guidance from L.J.F. and assisted B.B. with sample preparation and metabolomics data analysis. D.F.Q.S. performed all urease and cell wall laccase experiments and advised on ABTS antioxidant assays. J.W.K. and A.C. acquired funding for the project and L.J.F. funds the LSI Metabolomics Core Facility at UBC.

