## Extended Data Figures and Supplemental materials for "Glutathione metabolism impacts fungal virulence by modulating the redox environment"

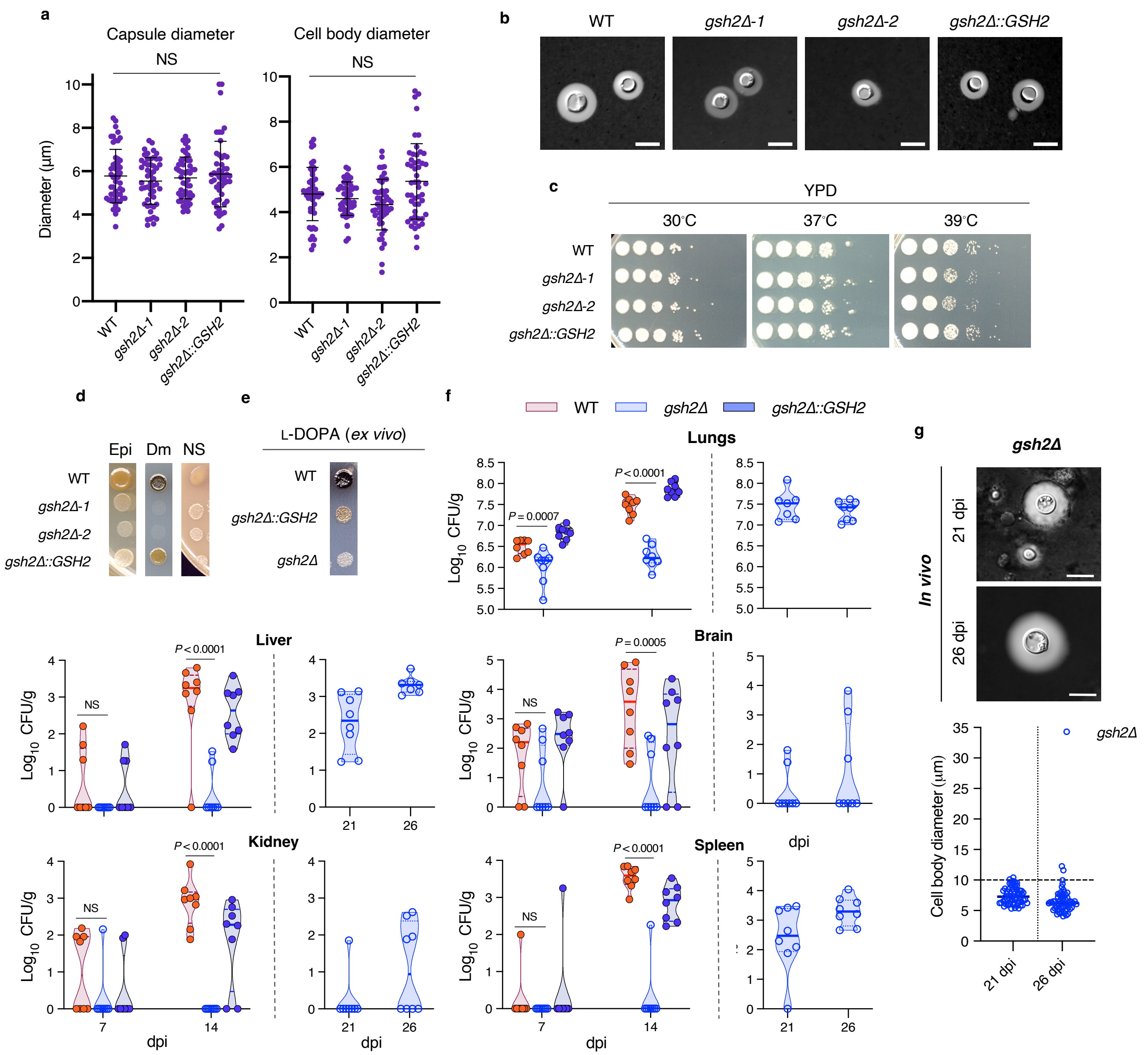


**Extended Data Figure 1. Loss of *GSH2* does not affect capsule size but delays virulence and impairs dissemination to the brain. a,** Polysaccharide capsule (left) and cell body (right) diameters of WT, *gsh2*Δ mutant, and *gsh2*Δ*::GSH2* strains grown for 48 h in CIM. Measurements represent mean values ± s.d. of 50 cells from each isolate with $n = 3$ biological replicates for each strain. **b,** Visualization of polysaccharide capsule with DIC microscopy.

Cells were negatively stained with 0.25 volumes of India ink. Images are representative of three independent experiments (scale bars = 10 μm). **c,** Spot assay of growth on solid YPD medium starting at $10^6$ cells $ml^{-1}$ with 10-fold serial dilutions. Plates were incubated at 30˚C, 37˚C, 39˚C. **d,** Melanin production on epinephrine (Epi, 0.1 g $L^{-1}$), dopamine (Dm, 0.1 g $L^{-1}$), and Niger seed (NS)-containing minimal medium. Each strain was plated on agar medium in triplicate at $10^6$ cells $ml^{-1}$ and grown for 48 h at 30°C before imaging. NS medium was prepared from 70 g 0.1 g $L^{-1}$ seed extract supplemented with 0.1 g $L^{-1}$ glucose, 20 g $L^{-1}$ Bacto Agar, and 0.05% Tween 20. All strains tested were unable to produce melanin on the indicated agar medium in the presence of exogenous GSH. **e,** Time-course of fungal burden in mice intranasally infected with WT (red), *gsh2*Δ (light blue), and *gsh2*Δ*::GSH2* (dark blue) strains. Groups of 8 mice were culled at each of the time points indicated for each strain. Note that the analysis was completed at day 14 for mice infected with the WT or complemented strains.  Statistical analysis of fungal burden was performed using a two-way ANOVA test with Dunnett's correction for multiple comparisons. Solid bars indicate mean fungal burden and segmented bars represent interquartile range. **f,** Visualization of fungal cells retrieved from murine lungs with DIC microscopy (top). Cells were negatively stained with 0.25 volumes of India ink. Images are representative of fungal cells retrieved from 8 murine lungs per strain ($n$ = 60 cells per sample) at each timepoint (bars = 10 μm). Cell body diameters of *gsh2*Δ mutants retrieved from murine lungs (bottom). Measurements represent mean values ± s.d. of 60 cells from each strain per time point from $n$ = 8 lungs. **g,** Melanin production of $10^6$ cells $ml^{-1}$ retrieved from murine and lungs spotted on solid L-DOPA medium for 48 h incubation at 30°C. Images are representative of three biological replicates

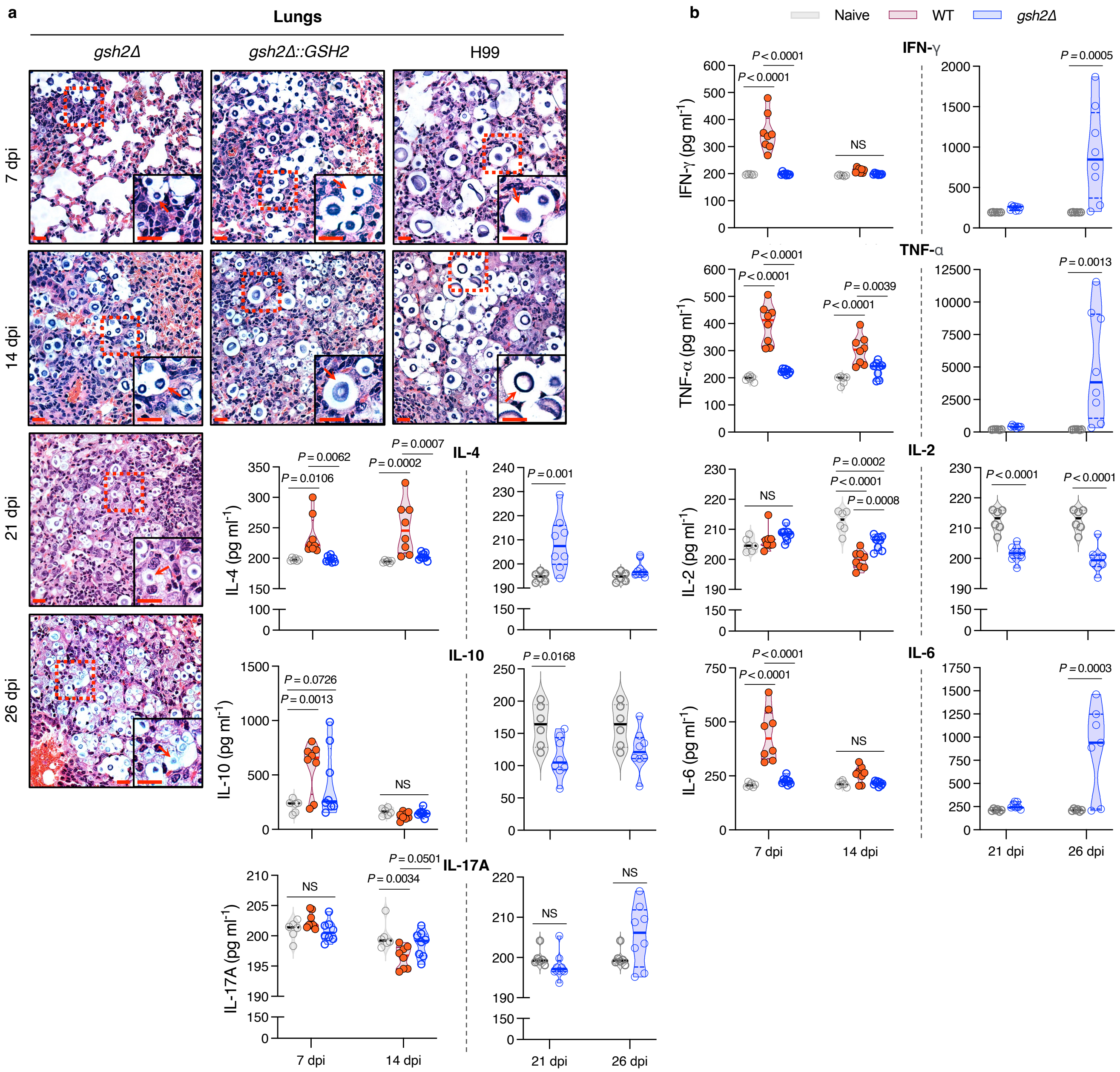


**Extended Data Figure 2. Mutants lacking GSH2 have reduced lung proliferation and cell size *in vivo*. a,** Representative images of H & E stained lung tissue sections from four mice infected with WT, *gsh2*Δ, or *gsh2*Δ*::GSH2* strains at each of the time points indicated (arrows = cryptococcal cells; scale bars = 10 μm). Segmented squares indicate enhanced zoom sections shown in the lower-right corner of each image. **b,** Cytokine levels in murine tissue homogenate of lungs infected with the WT or *gsh2*Δ mutant strains at the indicated timepoints (*n* = 8 lungs for each strain at each time point). Uninfected mice (Naïve, inoculated with PBS) were used as a control. Data were analyzed by two-way ANOVA with Tukey's correction for multiple comparisons. Solid bars indicate mean cytokine level and segmented bars represent interquartile range.

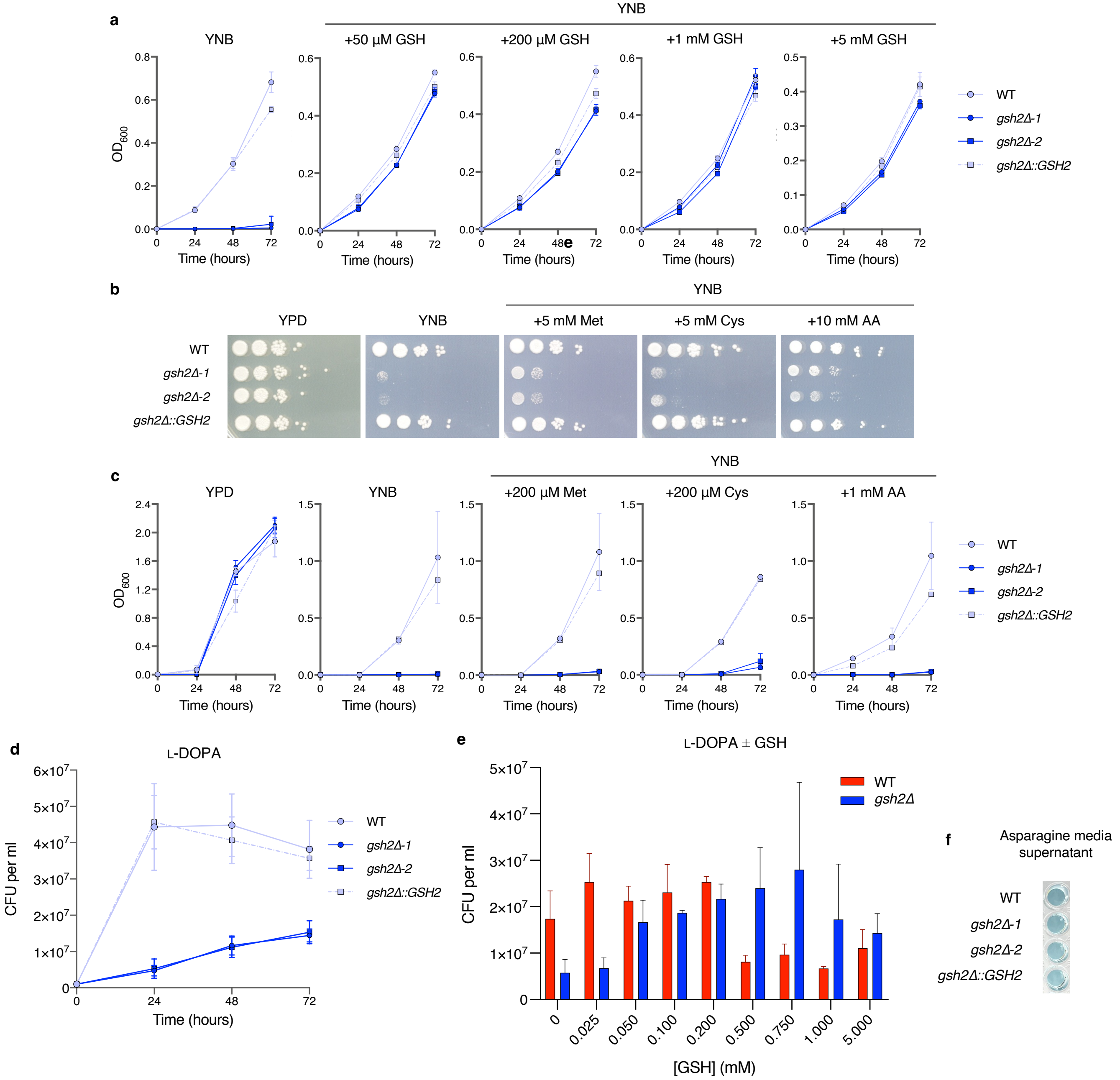


**Extended Data Figure 3. GSH is required for growth in minimal medium and is not restored with ascorbate or sulfur-containing amino acids. a,** Growth curve analysis of WT (light blue circles), *gsh2*Δ (blue squares or circles), and *gsh2*Δ*::GSH2* (light blue squares) strains grown in rich or minimal medium with and without methionine (Met), cysteine (Cys), or ascorbic acid (AA) supplementation at the indicated concentrations. **b,** Spot assays on minimal media. Each strain was spotted starting at $10^6$ cells $ml^{-1}$ with 10-fold serial dilutions on agar minimal medium with or without Met, Cys, or AA supplementation at the indicated concentrations. Images are representative of three biological replicates. **c,** Growth curve analysis of WT, *gsh2*Δ, and *gsh2*Δ*::GSH2* strains grown in minimal medium with or without GSH supplementation at the indicated concentrations. Data points for **a** and **c** indicate mean $OD_{600}$ values ± s.d., and the initial inoculum for each strain was $2 \times 10^4$ cells $ml^{-1}$. Growth was monitored for 72 h with $OD_{600}$ values measured every 24 h. **d–e,** Growth of WT, *gsh2*Δ, and *gsh2*Δ*::GSH2* cells grown in 5 ml liquid cultures of L-DOPA medium (with or without GSH supplementation in **e**). Initial inoculum for each strain was $10^6$ cells $ml^{-1}$ and CFUs were counted every 24 h for 72 h (**d**) or at 48 h (**e**). **f,** ABTS antioxidant assay (see Methods) for the proportion of ABTS radical quenched by supernatant isolated from the indicated strains after 72 h incubation in L-asparagine minimal medium (lacking L-DOPA). Pigmentation (blue colouration) indicates presence of the ABTS radical. Images represent three independent experiments. Data are representative of three biological replicates for each experiment.

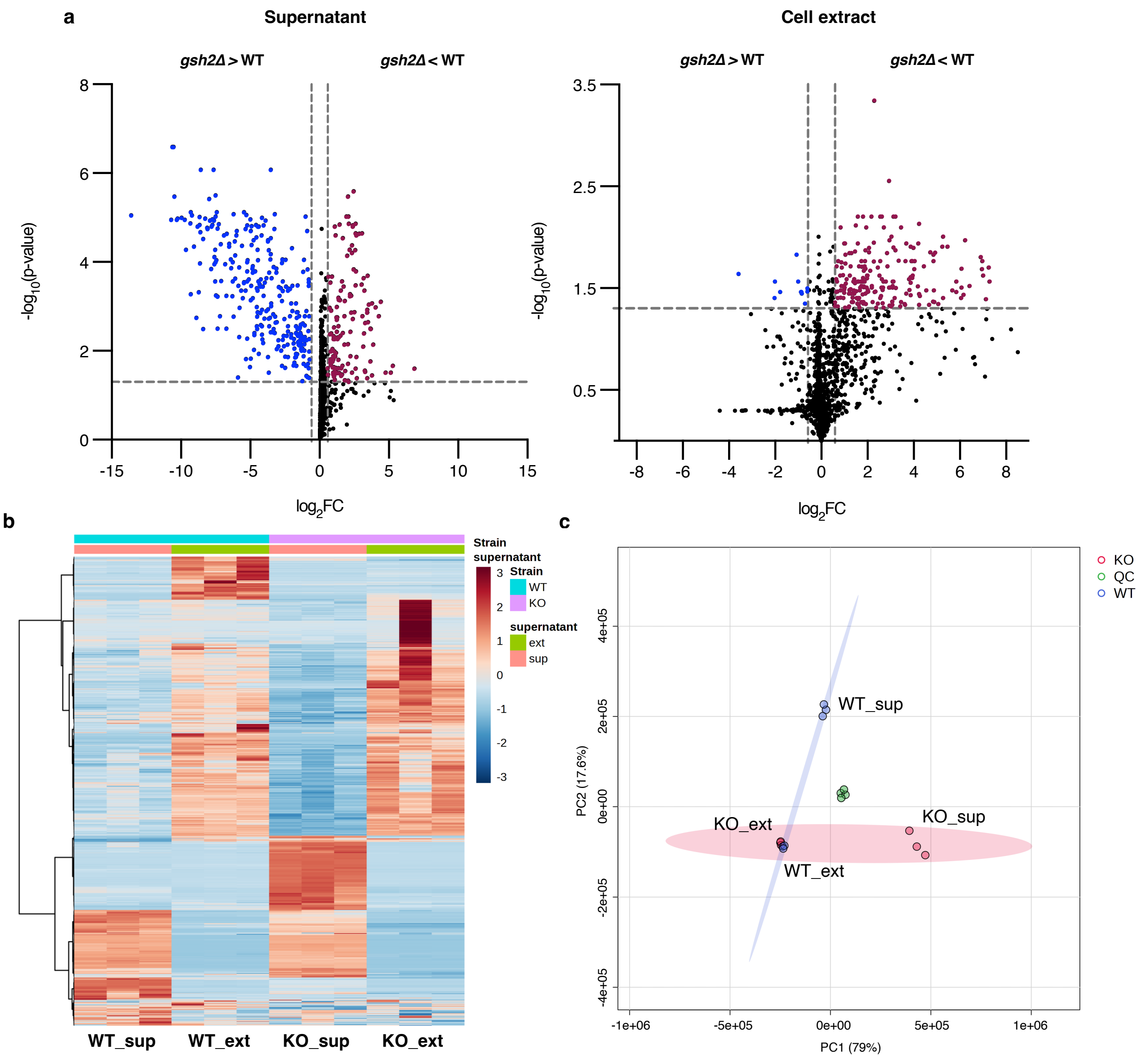


**Extended Data Figure 4. Metabolic profiling of the *gsh2*Δ mutant relative to WT reveals differences in extracellular and intracellular fractions. a,** Volcano plots showing QC-normalized LC-HRMS/MS feature data and differentially abundant metabolites between WT and *gsh2*Δ mutant cells in supernatant (left) and cell extract (right) fractions. The horizontal axis represents the directional intensity of the metabolite peak abundance fold change (FC) and the vertical axis represents statistical significance. A *p*-value threshold of $p < 0.05$ and FC threshold of $> 1.5$ or $< 0.667$ (segmented lines) were used to identify significant differences between the

WT and mutant strains, which were determined using an unpaired, two-tailed Student's *t*-test with FDR correction in MetaboAnalyst. **b,** Heatmap comparing relative intensity of metabolite abundances in the supernatant (sup) and cell extract (ext) of WT and *gsh2*Δ mutants (KO); $n$ = 3 biological replicates were analyzed for each fragment. Higher and lower intensity values are coloured red and blue, respectively. **c,** PCA score plot of the first two principal components from WT and *gsh2*Δ mutant supernatant and cellular extract fractions with QC samples for data normalization. Each data set represents $n$ = 3 biological replicates. Purple = WT; red = *gsh2*Δ mutant; green = QC.

**a**

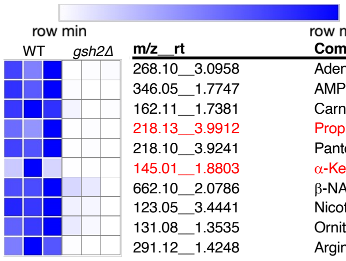


| m/z__rt | Compound | FC | log2FC | p.adjusted | -log10p.adj | IonMo | Frac |
|---|---|---|---|---|---|---|---|
| 268.10__3.0958 | Adenosine | 12.2217 | 3.611 | 0.045397 | 1.343 | pos | ext |
| 346.05__1.7747 | AMP | 14.5651 | 3.864 | 0.005449 | 2.264 | neg | ext |
| 162.11__1.7381 | Carnitine | 8.7303 | 3.126 | 0.012690 | 1.897 | pos | ext |
| 218.13__3.9912 | Propionylcarnitine | 20.7093 | 4.372 | 0.080483 | 1.094 | pos | ext |
| 218.10__3.9241 | Pantothenate | 37.2877 | 5.221 | 0.011592 | 1.936 | neg | ext |
| 145.01__1.8803 | α-Ketoglutaric acid | 11.3842 | 3.509 | 0.297301 | 0.527 | neg | ext |
| 662.10__2.0786 | β-NAD | 2.6050 | 1.381 | 0.021111 | 1.675 | neg | ext |
| 123.05__3.4441 | Nicotinamide | 2.9898 | 1.580 | 0.011530 | 1.938 | pos | ext |
| 131.08__1.3535 | Ornithine | 3.0519 | 1.610 | 0.015763 | 1.802 | neg | ext |
| 291.12__1.4248 | Arginosuccinic acid | 25.1902 | 4.655 | 0.046401 | 1.333 | pos | ext |

**b**

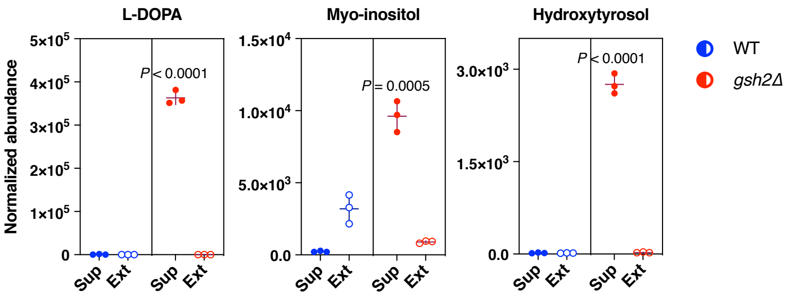


**c**

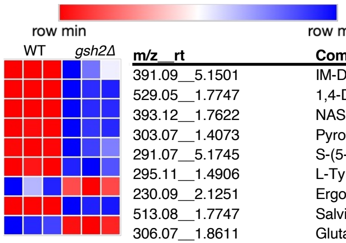


| m/z__rt | Compound | FC | log2FC | p.adjusted | -log10p.adj | IonMo | Frac |
|---|---|---|---|---|---|---|---|
| 391.09__5.1501 | IM-DSGLS | 0.0253 | -5.307 | 0.015530 | 1.809 | pos | sup |
| 529.05__1.7747 | 1,4-DMGLB | 0.0075 | -7.051 | 0.000053 | 4.272 | neg | sup |
| 393.12__1.7622 | NAS glucuronide | 0.0009 | -10.053 | 0.000057 | 4.245 | neg | sup |
| 303.07__1.4073 | Pyrogallol-2-O-glucuronide | 0.0131 | -6.258 | 0.000104 | 3.984 | pos | sup |
| 291.07__5.1745 | S-(5-his)cys SO | 0.1080 | -3.211 | 0.001753 | 2.756 | pos | sup |
| 295.11__1.4906 | L-Tyrosyl-L-methionine | 0.0196 | -5.672 | 0.000452 | 3.345 | pos | sup |
| 230.09__2.1251 | Ergothioneine | 2.2503 | 1.170 | 0.011352 | 1.945 | pos | sup |
| 513.08__1.7747 | Salvianolic acid C | 0.0143 | -6.128 | 0.000027 | 4.576 | neg | sup |
| 306.07__1.8611 | Glutathione (reduced) | 5.5313 | 2.468 | 0.000208 | 3.682 | neg | sup |

**d**

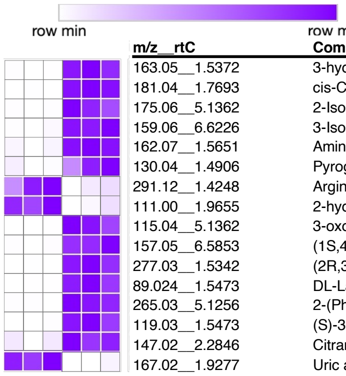


| m/z__rtC | Compound | FC | log2FC | p.adjusted | -log10p.adj | IonMo | Frac |
|---|---|---|---|---|---|---|---|
| 163.05__1.5372 | 3-hydroxy-3-MGA | 0.0522 | -4.259 | 0.000365 | 3.437 | pos | sup |
| 181.04__1.7693 | cis-Caffeic acid | 0.0157 | -5.998 | 0.000026 | 4.584 | pos | sup |
| 175.06__5.1362 | 2-Isopropylmalic acid | 0.0338 | -4.888 | 0.000911 | 3.040 | neg | sup |
| 159.06__6.6226 | 3-Isopropylmalic acid | 0.1812 | -2.464 | 0.000092 | 4.036 | pos | sup |
| 162.07__1.5651 | Aminoadipic acid | 0.2502 | -1.999 | 0.000051 | 4.293 | pos | sup |
| 130.04__1.4906 | Pyroglutamic acid | 0.4830 | -1.050 | 0.028882 | 1.539 | pos | sup |
| 291.12__1.4248 | Argininosuccinic acid | 2.9578 | 1.565 | 0.036930 | 1.433 | pos | sup |
| 111.00__1.9655 | 2-hydroxyGA lactone | 1.8915 | 0.920 | 0.002391 | 2.621 | neg | sup |
| 115.04__5.1362 | 3-oxovaleric acid | 0.0438 | -4.512 | 0.001388 | 2.858 | neg | sup |
| 157.05__6.5853 | (1S,4S)-4-hydroxy-3-OCHCA | 0.0511 | -4.291 | 0.000549 | 3.260 | neg | sup |
| 277.03__1.5342 | (2R,3S)-Piscidic acid | 0.0159 | -5.972 | 0.000343 | 3.465 | neg | sup |
| 89.024__1.5473 | DL-Lactic Acid | 0.0241 | -5.375 | 0.000642 | 3.192 | neg | sup |
| 265.03__5.1256 | 2-(Phenylthio)BA | 0.0492 | -4.347 | 0.000175 | 3.757 | neg | sup |
| 119.03__1.5473 | (S)-3,4-dihydroxybutyric acid | 0.0577 | -4.116 | 0.000773 | 3.112 | neg | sup |
| 147.02__2.2846 | Citramalic acid | 0.2802 | -1.835 | 0.000923 | 3.035 | neg | sup |
| 167.02__1.9277 | Uric acid | 8.7751 | 3.133 | 0.000853 | 3.069 | neg | sup |

**Extended Data Figure 5. Mutants lacking *GSH2* show distinct changes in relative abundance of key energy metabolites, antioxidants, and extracellular acids. a–d**, Relative abundance of compounds identified via LC-HRMS/MS between WT (blue) and *gsh2*Δ mutant (red) strains. Both cell extract (ext) and supernatant (sup) fractions were analyzed for relative peak intensity. **a,** Energy pathway and urea cycle intermediates are depleted in the cellular extract of *gsh2*Δ mutants. AMP = adenosine monophosphate; β-NAD = β-nicotinamide adenine dinucleotide. Significance was determined via a one-way ANOVA with FDR correction. **b,** L-DOPA is over 500-fold more abundant in the *gsh2*Δ mutant supernatant compared to the WT strain. The anti-melanogenic compounds myo-inositol and hydroxytyrosol are also highly abundant in the *gsh2*Δ mutant supernatant. Bars represent the mean relative abundance ± s.d. for $n = 3$ biological replicates per strain. Statistical significance was calculated relative to the WT supernatant (sup) fraction using a one-way ANOVA with FDR correction. **c,** Compounds with known antioxidant activity are highly abundant in the *gsh2*Δ mutant supernatant fraction compared to WT (except ergothioneine), suggesting an extracellular reducing environment of mutant strains. IM-DSGLS = indolylmethyl-desulfoglucosinolate; 1,4-DMGLB = 1,4-dimethoxcyglucobrassicin; NAS glucuronide = N-acetylserotonin glucuronide; S-(5-his)cys SO = S-(5-histidyl)cysteine sulfoxide. **d,** *gsh2*Δ mutants accumulate ketone bodies and various acids and acid derivatives in the supernatant that may contribute to low pH and an extracellular reducing environment. 3-hydroxy-3-MGA = 3-hydroxy-3-methyl-Glutaric acid; 2-hydroxyGA lactone = 2-hydroxy-Glutaric acid lactone; (1S,4S)-4-hydroxy-3-OCHCA = (1S,4S)-4-hydroxy-3-oxocyclohexanecarboxylic acid; 2-(Phenylthio)BA = 2-(Phenylthio)benzeneacetic acid. All heat maps were generated using Morpheus matrix visualization and analysis software (https://software.broadinstitute.org/morpheus/). Compounds in red text were not statistically significant after FDR correction but had high FC values.

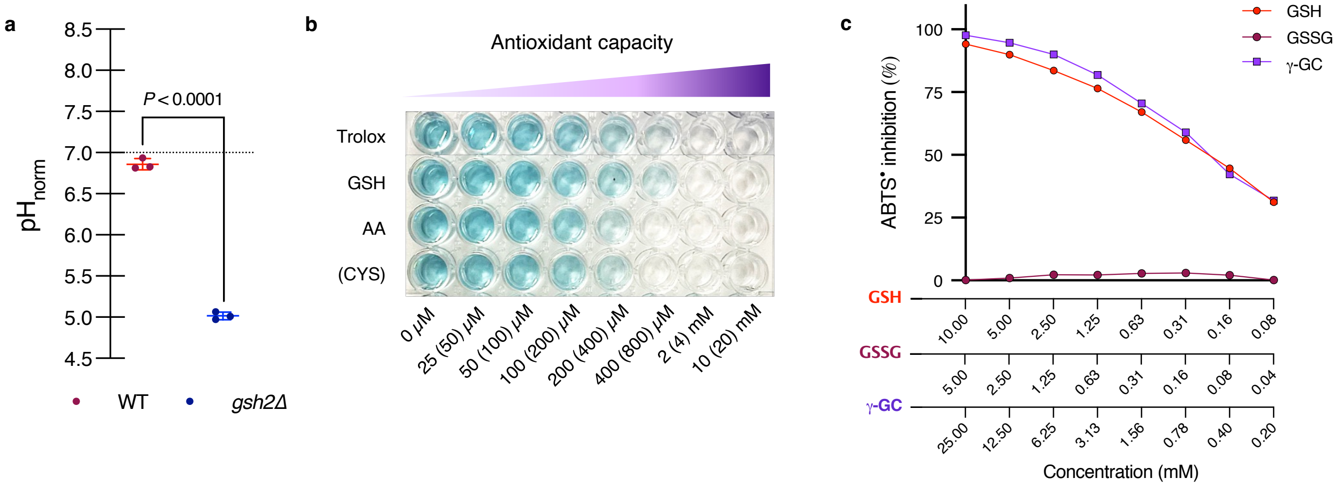


**Extended Data Figure 6. Extracellular acidification of *gsh2*Δ mutants is independent of L-DOPA and GSH pathway metabolites influence melanin formation. a,** pH values of supernatant isolates from WT and *gsh2*Δ mutant cells grown in L-asparagine minimal medium and normalized to $10^7$ cells $ml^{-1}$. Statistical significance was calculated using an unpaired, two-tailed Student's *t*-test. Bars represent mean pH values ± s.d. **b–c,** ABTS antioxidant assay indicates the reducing power of compounds tested at the indicated concentrations. Decreased pigmentation (light blue & clear in image) in **c** indicates increased $ABTS^{\bullet}$ radical scavenging activity. GSH = glutathione; AA = ascorbic acid; Cys = cysteine. **d,** Melanin production of liquid cultures incubated for 72 h in L-DOPA medium with or without 5 mM GSH, 2.5 mM GSSG, or 5 mM γ-glutamylcysteine (γ-Glu-Cys). Absorbance of cell supernatant at $OD_{490}$ was measured before (left, extracellular melanin) and after (right, CWB melanin) digestion with 1 M NaOH + 10% DMSO for 1 h at 95°C and normalized to $OD_{600}$. Bars represent the mean ± s.d. Significance relative to WT was calculated using two-way ANOVA with Dunnett's correction for multiple comparisons. Data are representative of three biological replicates for each experiment.

### Supplementary Figures

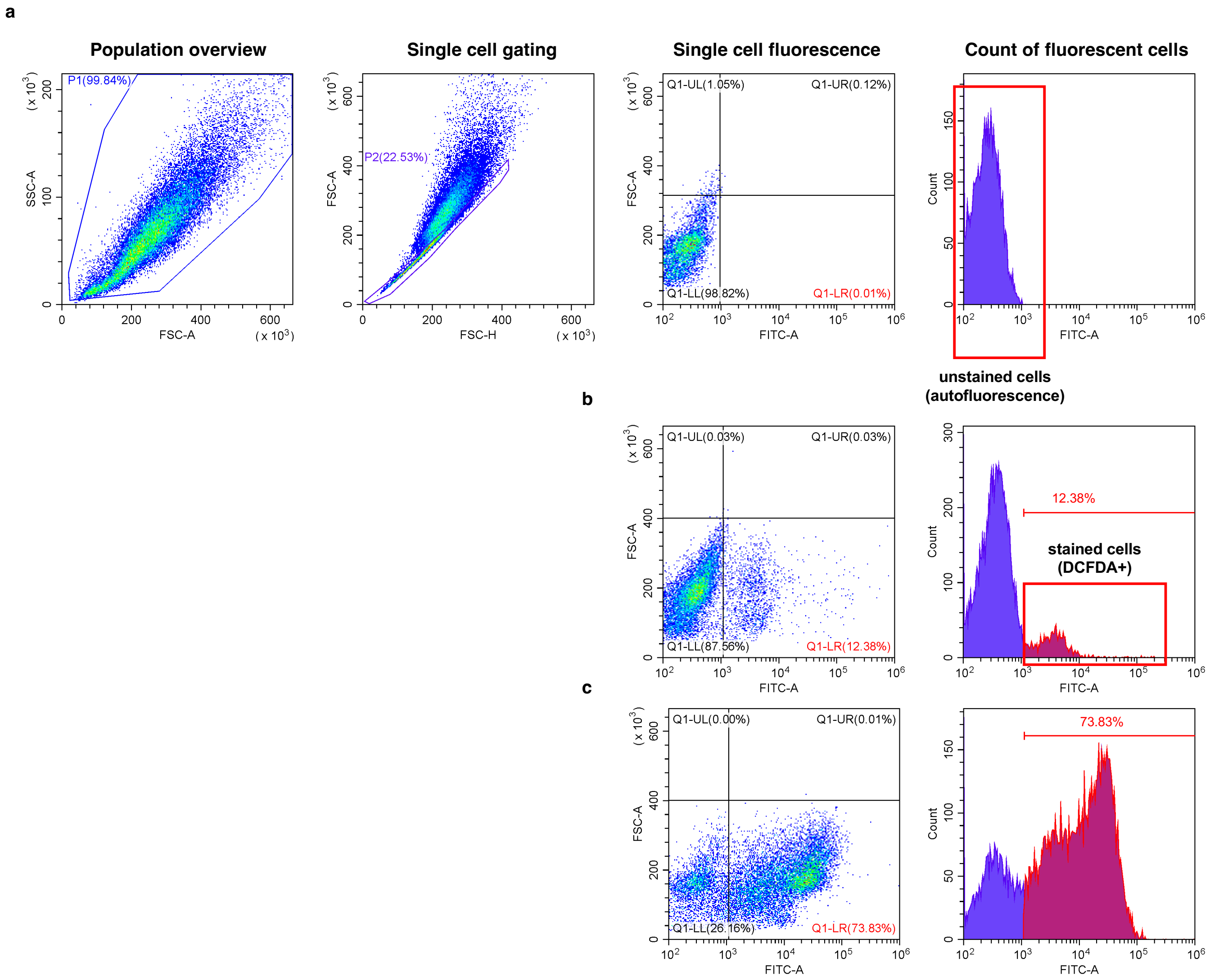


**Supplementary Figure 1. Gating strategy for ROS-accumulation assay using flow cytometry. a,** Gating strategy for flow cytometry first identifies all *C. neoformans* cells (P1, left) detected by forward (FSC-A) and side (SSC-A) scatter area parameters. Single cells (P2) were further delineated based on area (FSC-A) and height (FSC-H) parameters to exclude cell doublets, and analyzed for fluorescence using the FITC-GFP (FITC-A, right) channel. Cells depicted in **a** are unstained and used as a control. **b–c,** Cells stained with the fluorogenic probe 2’,7’-dichlorodihydrofluorescein diacetate (DCFDA) and separated based on the parameters outlined in **a**. Cells were either untreated (**b**) or treated (**c**) with 1 mM $H_2O_2$ prior to DCFDA staining to induce ROS accumulation.

#### Supplementary Tables

**Table S1. List of *Cryptococcus neoformans* strains.**

| Strain | Description |
|---|---|
| H99 | Wild-type (WT) strain and background for mutants generated for this study. |
| *gsh2-1*Δ | Independent gene knockout mutant lacking *GSH2* |
| *gsh2-2*Δ | Independent gene knockout mutant lacking *GSH2* |
| *gsh2*Δ*::GSH2* | Strain complemented with WT *GSH2* in the *gsh2*Δ mutant background |

**Table S2. List of primers used for strain construction.**

Primers for strain construction:

| Strain construct | Primer name | Sequence (5' – 3') |
|---|---|---|
| *gsh2*Δ mutant | Gsh2-1 | GGGTGCAGCTAATGTCACAA |
| | Gsh2-2 | CATACATCCTCTTCTTCCCCACCGCTGCGAGGATGTGAGCT |
| | Gsh2-3 | AGCTCACATCCTCGCAGCGGTGGGGAAGAAGAGGATGTATG |
| | Gsh2-4 | TAGTTTCTACATCTCTTCTACGCGAACTGCTTCTTCTGTGTC |
| | Gsh2-5 | GACACAGAAGAAGCAGTTCGCGTAGAAGAGATGTAGAAACTA |
| | Gsh2-6 | GCCAAGCAACAAGCAATGTATGA |
| | Gsh2-9PO | GCTCTTTCTGGATCCGATCTAGT |
| | Gsh2-10PO | CTGAACATCACTGTAAGGTGCGTC |
| *gsh2*Δ*::GSH2* complement | Gsh2_HA1 | TCCTCTTCGAGCAGTGTTGC |
| | Gsh2_HA2 | AGGGACATCGTAAGGGTAATCCACAAGCAAAGGACTGTCA |
| | Gsh2_HA3 | GTCCTTTGCTTGTGGATTACCCTTACGATGTCCCTGATTACG |
| | Gsh2_HA4 | GTTCGCGTATGCATAAGCAGATGTAGAAACTAGCTTCCTGG |
| | Gsh2_HA5 | GGAAGCTAGTTTCTACATCTGCTTATGCATACGCGAACTGC |
| | Gsh2_HA6 | CTGCTGAATCAGTGGCTTTCG |
